# Analysis of calcium dynamics for dim-light responses in rod and cone photoreceptors

**DOI:** 10.1101/2022.03.22.485340

**Authors:** Annia Abtout, Jürgen Reingruber

## Abstract

Rod and cone photoreceptors in the retina of vertebrate eyes are fundamental sensory neurons underlying vision. They use a sophisticated signal transduction pathway consisting of a series of biochemical processes to convert the absorption of light into an electrical current response. Several of these processes are modulated by feedback that depends on the intracellular *Ca*^2+^ concentration. In this work we use a representative phototransduction model to study how changing the *Ca*^2+^ kinetics by fast buffering affects sensitivity and dynamics of the light response in mouse rod and cone photoreceptors. We derive analytic results for dim-light stimulations that provide quantitative and conceptual insight. We show that flash responses are monophasic with low buffering, and the change in the *Ca*^2+^ concentration occurs in proportion to the current. If the amount of fast buffering is increased, the *Ca*^2+^ kinetics becomes slowed down and delayed with respect to the current, and biphasic responses emerge (damped oscillations). This shows that a biphasic response is not necessarily a manifestation of slow buffering reactions. A phase space analysis shows that the emergence of biphasic responses depends on the ratio between the effective rate *μ*_*ca*_ that controls the *Ca*^2+^ kinetics, and the dark turnover rate of cyclic GMP *β*_*d*_. We further investigate how the light response is altered by modifying the extracellular *Ca*^2+^ concentration. In summary, we provide a comprehensive quantitative analysis that precisely links the dynamics of *Ca*^2+^ concentration to the observed current response.

## 1 Introduction

Vision in most vertebrates is sustained by rod and cone photoreceptors in the retina of the eyes [1]. Whereas rods are most sensitive to light and sustain vision in dim-light conditions, cones have a much reduced light sensitivity, but they adapt and thereby maintain vision even under brightest light conditions [2].

The signal transduction cascade in the outer segment of rod and cone photoreceptors is complex multistep process that transforms the absorption of light by photopigments into an electrical current (for reviews see [3, 4, 5, 6]). Many of these processes are modulated by *Ca*^2+^ feedback [7, 8, 9, 10, 11]. *Ca*^2+^ feedback not only controls how photoreceptors adapt to increasing light intensities [12, 13, 14, 15, 16], it also shapes the response in darkness [17, 18, 19, 20]. Dark adapted rods in most species show monophasic responses to brief flashes of light (monophasic means that the current is always reduced compared to its steady state level in darkness). If the *Ca*^2+^ kinetics is disturbed by the application of exogenous buffers, biphasic responses emerge (biphasic means that the current also reaches values that are larger than the steady state level). With exogenous *Ca*^2+^ buffers, biphasic flash responses have been observed in amphibian rods [21, 22, 20, 23, 24], in primate and guinea pig rods [25, 17, 14], and in mouse rods [17, 26]. In primate and mammalian cones, biphasic responses have often been observed even without exogenous buffer application [27, 28, 29, 30, 31, 32]. However, recently it has been claimed that cone responses are mostly monophasic under physiological conditions, and that biphasic responses are most likely the result of a distorted negative *Ca*^2+^ feedback [33]. There it has also been shown that biphasic cone responses could be removed by lowering the extracellular *Ca*^2+^ concentration [33].

The dynamics of the free-*Ca*^2+^ concentration in the OS depends on influx through cyclic nucleotide-gated (CNG) channels, efflux via NCKX exchangers, and cytoplasmic buffering [6]. With negligible *Ca*^2+^ buffering, modelling shows that the *Ca*^2+^ concentration changes in proportion to the current, and flash responses are monophasic [34, 35]. Biphasic responses have been simulated with models that take into account the effect of buffering [36]. One possibility to model the effect of buffering is to consider explicit equations for the binding between free *Ca*^2+^ and the buffers [36, 37, 38, 39, 40, 41, 42]. A different approach, which is more frequently used, is to lump the effect of the various buffers into a single parameter called the buffering capacity *B*_*ca*_ [43, 44, 6, 45, 46, 47, 48, 49, 50] (a *Ca*^2+^-dependent buffering capacity has been used in [27]). The use of a constant buffering capacity is a valid approximation if the binding kinetics are fast (fast buffers), and the *Ca*^2+^ affinity of the buffers is low compared to the physiological range of the *Ca*^2+^ concentration. Thus, whereas modelling of the binding reactions between *Ca*^2+^ and buffers is most accurate, models with a buffering capacity are an approximation, but simpler to handle.

So far, insight from modelling about the *Ca*^2+^ dynamics and the effect of buffering has been obtained almost exclusively from numerical simulations (but see the analysis in [43, 30, 44, 28]). Recently, we derived analytic results for the entire dim-light response that provided new insight about how the various biophysical processes determine amplitude and shape of flash responses in rods and cones [34, 35]. In this previous work, we neglected the effect of buffering, and assumed that the free-*Ca*^2+^ concentration changes in proportion to the current, which often is the physiological case [51, 19]. In the present manuscript, we generalize our previous analysis by studying how the dim-light response is modified if the *Ca*^2+^ dynamics becomes gradually delayed with respect to the current due to buffers with fast binding kinetics. We derive analytic formulas that reproduce a dim-light response in presence of buffering, which we use to obtain quantitative and conceptual insight. We show that biphasic responses (damped oscillations) emerge as the *Ca*^2+^ dynamics is slowed down. We study the phase space that separates monophasic from biphasic responses, and we determine how sensitivity and shape of responses are affected by buffering. Finally, we examine the impact of changing the extracellular *Ca*^2+^ concentration.

## 2 Materials and Methods

### 2.1 Phototransduction model

We start from the parsimonious phototransduction model introduced in [35, 34], and we include a more general *Ca*^2+^ dynamics where the free-*Ca*^2+^ concentration does not necessarily change in proportion to the circulating current. In short, light activates the visual pigment *R*^*∗*^ with a rate that is proportional to the light intensity *ϕ*(*t*) times the collecting area *κ. R*^*∗*^ activates the G-protein transducin *T*^*∗*^ with a rate *k*_*act*_, and deactivates with rate *μ*_*rh*_. *T*^*∗*^ activates phosphodiesterase (PDE) *P*^*∗*^ with a rate *k*_*tr*_, and deactivates with rate *mu*_*tr*_. Because the deactivation of *T*^*∗*^ is linked to the activation of PDE, we have *k*_*tr*_ = *μ*_*tr*_ (to keep the analysis most general, we distinguish between *k*_*tr*_ and *μ*_*tr*_). *P*^*∗*^ deactivates with rate *μ*_*pde*_, and hydrolyses the cytosolic second messenger cyclic GMP (cGMP) with rate constant *β*_*sub*_. cGMP gates the opening of CNG channels, in the OS membrane. The rate *α* by which the cGMP concentration *c*_*cg*_ is synthesized by guanylate cyclase (GC) depends on the free-*Ca*^2+^ concentration 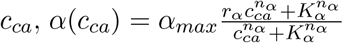, where 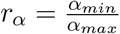. The cyclase activity is modulated via the intermediate binding of *Ca*^2+^ to GCAP proteins [52, 17]. *Ca*^2+^ also affects the phosphorylation dependent deactivation of the photopigment by binding to recoverin, which inhibits the photopgiment phosphorylation by rhodopsin kinase [53, 54]. However, in this work we only consider the *Ca*^2+^ feedback to guanylate cyclase, which is most important under dim light conditions [17, 18, 19].

The *Ca*^2+^ concentration in the OS changes due to influx via CNG channels, and efflux via electrogenic NCKX exchangers. The CNG current is *I*_*ch*_ = *I*_*ch,max*_*p*_*ch*_, where 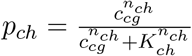 is the cGMP dependent fraction of open channels, and *K*_*ch*_ is the *cGMP* concentration where 50% of the channels are open. Only the fraction *f*_*ch,ca*_ of the CNG current is carried by *Ca*^2+^, *I*_*ch,ca*_ = *f*_*ch,ca*_*I*_*ch*_. The influx of *Ca*^2+^ via CNG channels therefore is 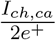. The exchanger current as a function of the *Ca*^2+^ concentration is *I*_*ex*_ = *I*_*ex,sat*_*p*_*ex*_, where *I*_*ex,sat*_ is the saturating current at high *c*_*ca*_, and 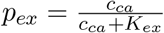 is the saturation level of the exchangers. The exchanger stoichiometry is such that the extrusion of a single *Ca*^2+^ leads to the influx of a single positive charge *e*^+^, such that *I*_*ex*_*/e*^+^ gives the *Ca*^2+^ efflux.

*Ca*^2+^ binds to and regulates the activity of many proteins participating in the signal transduction cascade [15, 9, 10, 11]. In principle, all these reactions contribute to *Ca*^2+^ buffering. With *N*_*b*_ buffer species *b*_*i*_ (*i* = 1, … *N*_*b*_), the *Ca*^2+^ dynamics in the OS is governed by the equations

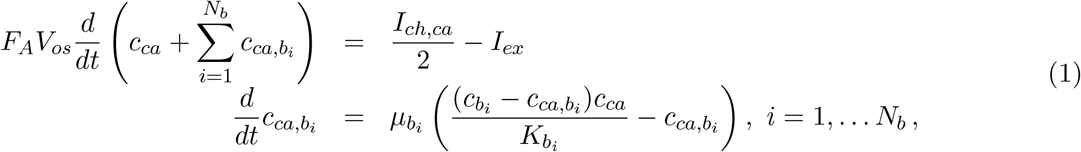

where 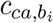 are the *Ca*^2+^ concentrations that are bound to the buffer species 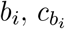 are total buffer concentrations, 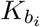 are dissociation constants, 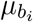 are dissociation rates, 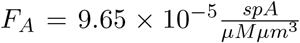 is the Faraday constant, and *V*_*os*_ is the outer segment volume. In this work we restrict the analysis to buffering reactions that proceed much faster compared to the time scale of the current change. With fast buffering (large 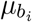), 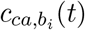 becomes a function of the free-*Ca*^2+^ concentration 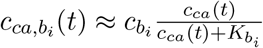 (quasi steady-state approximation). With fast buffers, Eq. 1 simplifies to

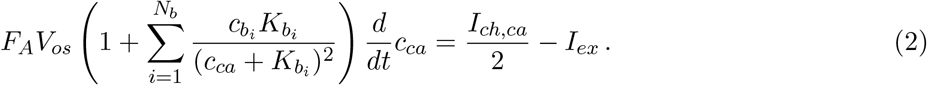

The complete system of transduction equations reads

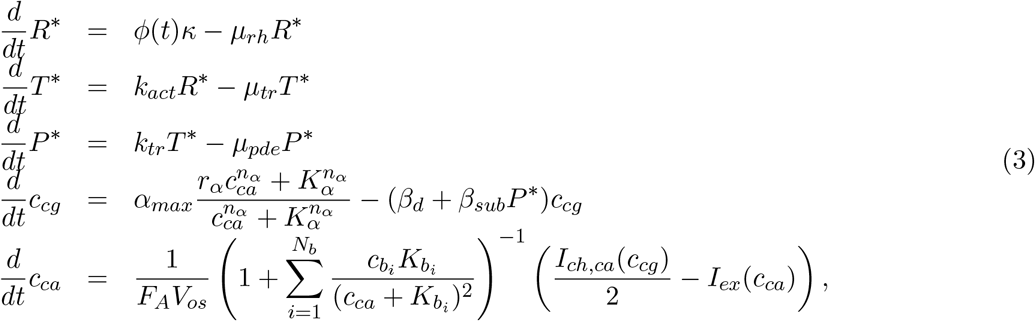

where *β*_*d*_ is the basal cGMP hydrolysis rate in darkness. The total current is

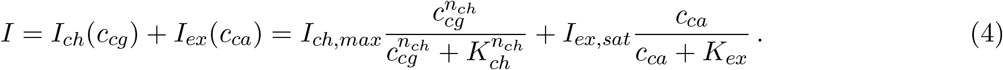

Next we use the concentrations *c*_*ca*,0_ and *c*_*cg*,0_, and the current *I*_0_ to introduce normalized variables and parameters. Since we are interested in the light response in darkness, we choose *c*_*ca*,0_, *c*_*cg*,0_, and *I*_0_ to be steady state values in darkness. These values can either be computed from Eq. 3 with *ϕ*(*t*) = 0, or one can directly use the experimentally measured values for *c*_*ca*,0_, *c*_*cg*,0_ and *I*_0_. In the latter case, we implicitly replaces the parameters *α*_*max*_, *I*_*ch,max*_ and *I*_*ex,sat*_ (which are not well known) with *c*_*ca*,0_, and *I*_0_. We introduce the normalized variable and parameters 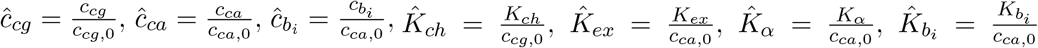, and we define the dark buffering constants 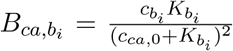, such that 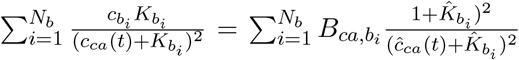. Because *c*_*ca*,0_ and *c*_*cg*_,_0_ are steady state values, we have *α*(*c*_*ca*,0_) = *β*_*d*_*c*_*cg*,0_ and *I*_*ch,ca*_(*c*_*cg*,0_) = *f*_*ch,ca*_*I*_*ch*_(*c*_*cg*,0_) = 2*I*_*ex*_(*c*_*ca*,0_). To simplify the analysis, we further introduce 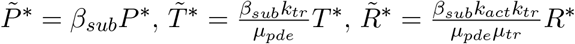, and the gain 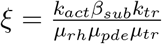 [35]. With the steady state condition *α*(*c*_*ca*,0_) = *β*_*d*_*c*_*cg*,0_ we obtain from Eq. 3

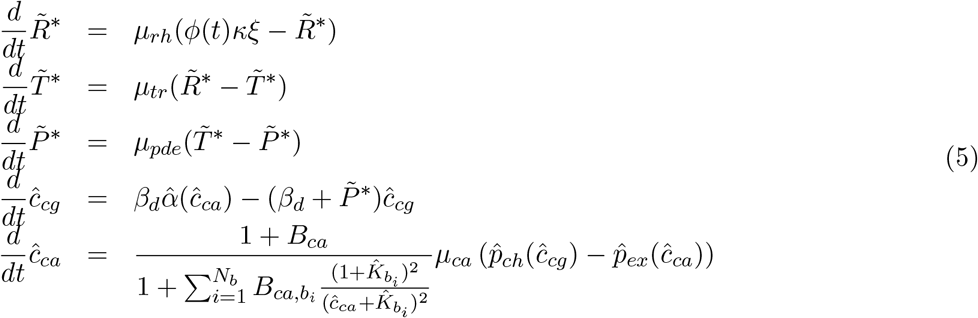

with

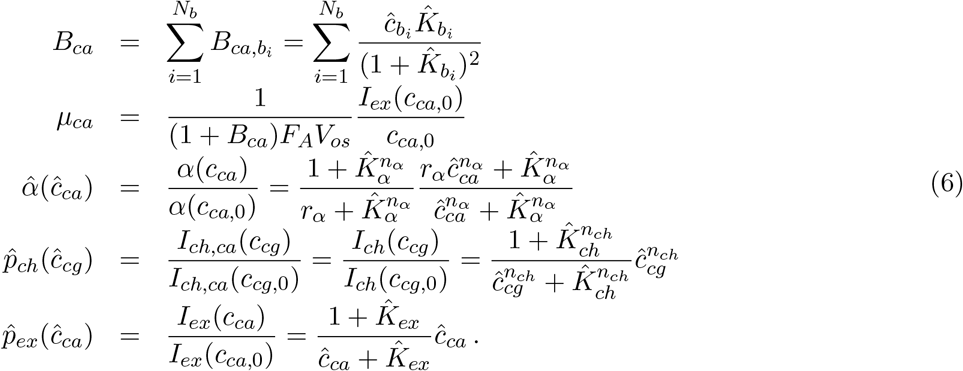

The conditions in darkness with *ϕ* = 0 are 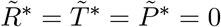 and 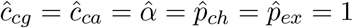.

The current in Eq. 4 normalised by the dark current 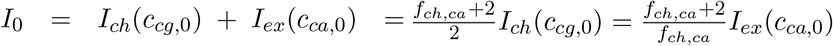 is

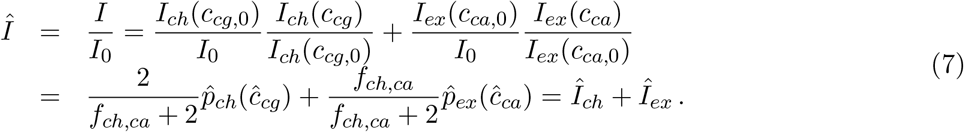

In darkness we have *Î* = 1. We further introduce the normalized current

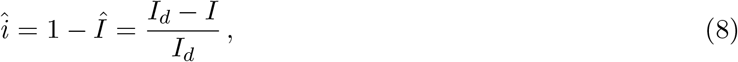

which is zero in darkness. 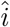 is usually used in the literature to characterize the light response. Although we refer to 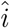 as a current (as it is usually done), it is clear that 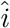 represents the current change with respect to the dark current. *Î* and 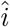 have complementary behaviours, for example, whereas *Î* decreases after the light is switched on, 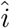 increases.

Before starting with the analysis of Eq. 5, we add some clarifying remarks:

1. Because we study the light response in darkness, we define the buffering constants 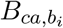 using the dark *Ca*^2+^ concentration. With a constant background light, we would define the 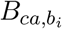 using the *Ca*^2+^ concentration corresponding to this background light. With high affinity buffers, this will adapt the value of the buffering capacity *B*_*ca*_ to the lower *Ca*^2+^ range in presence of a light background.
2. Per definition, 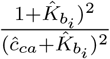 is one in darkness where *ĉ*_*ca*_ = 1. As we will show below, in first order it can be approximated by one for the whole dim-light response. Hence, for dim light we have 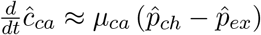.
3. Eq. 3 and Eq. 5 are mathematically equivalent, however, Eq. 5 depends on fewer parameters, which simplifies analysis and comprehension. Eq. 5 depends only on parameters that are important for the response dynamics. For example, Eq. 5 reveals that the activation rates *k*_*act*_, *k*_*tr*_ and *β*_*sub*_ affect the light response only as the product *k*_*act*_*k*_*tr*_*β*_*sub*_. Moreover, Eq. 5 shows that the *Ca*^2+^ dynamics is determined by the effective rate *μ*_*ca*_.
4. Eq. 5 is derived assuming that the normalisation values *c*_*ca*,0_ and *c*_*cg*,0_ are the correct steady state concentrations in darkness 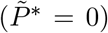. Eq. 5 can be used to study the response dynamics as a function of the light intensity. In contrast, if one wants to study how the steady state is altered when parameter values are changed, Eq. 5 has to be adapted. For example, if the cyclase sensitivity is altered from 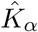 to a new value 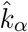, the synthesis rate *α*(*c*_*ca*,0_) changes from *α*(*c*_*ca*,0_) = *β*_*d*_*c*_*cg*,0_ to *α*(*c*_*ca*,0_) = *ζβ*_*d*_*c*_*cg*,0_, where 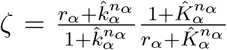. The correct equation for *ĉ*_*ca*_ is 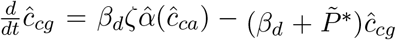, with 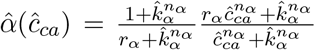. It follows that *ĉ*= *ĉ* = 1 are no longer steady state solutions in darkness. The new steady values *ĉ*_*ca*_ and 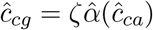 are obtained by solving the equation 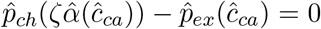. In a subsequent step, one can use the new steady state values to renormalise parameters, in which case one again obtains Eqs. 5,6, but with altered normalized parameters (see the analysis with modified extracellular *Ca*^2+^ below).

### 2.2 Linear response analysis for dim-light stimulations

PDE activation in Eq. 5 is linear and can be solved analytically,

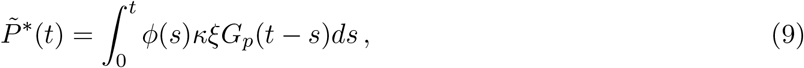

with the Green’s function

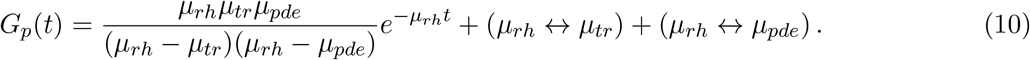

The double-headed arrows in Eq. 10 signify that the second and third terms in the equation are identical to the first term with the exchange *μ*_*tr*_ and *μ*_*rh*_ in the second term, and *μ*_*pde*_ and *μ*_*rh*_ in the third term. The equations for *ĉ*_*cg*_ and *ĉ*_*ca*_ (Eq. 5) cannot be solved analytically for all conditions of light stimulation *ϕ*(*t*). However, analytic results can be derived for dim-light intensities by linearizing Eq. 5. For the analysis we introduce *y*(*t*) = − ln *ĉ*_*cg*_(*t*) and *z*(*t*) = − ln *ĉ*_*ca*_(*t*), which are zero in darkness. For dim-light responses with 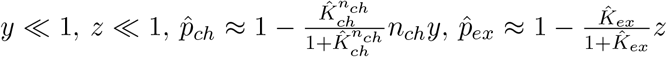, we obtain from Eq. 5 the first-order equations

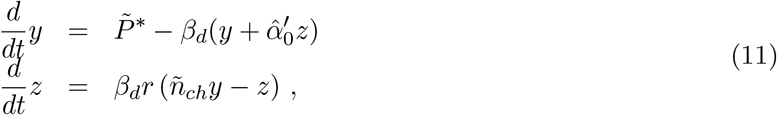

where

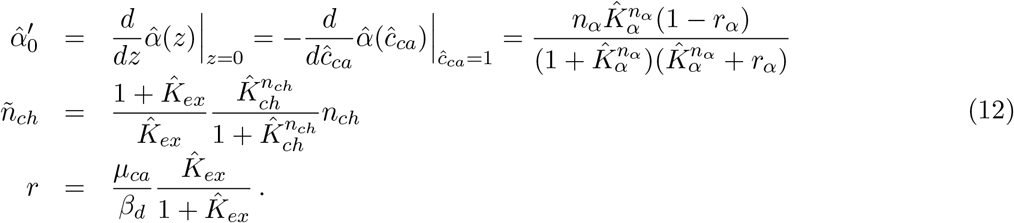

For GCAPs^−/−^ mutants the cyclase is not *Ca*^2+^ dependent and we have 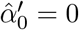. The two independent solutions of Eq. 11 with 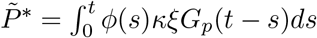 are

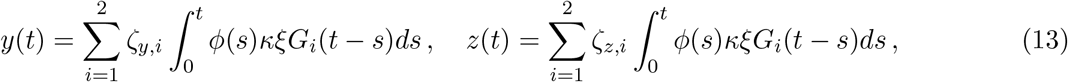

where 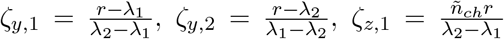 and 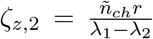. The parameters 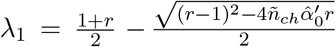 and 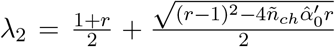 are the eigenvalues of the matrix 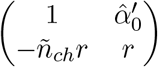.

The two independent Green’s functions are (*i* = 1, 2)

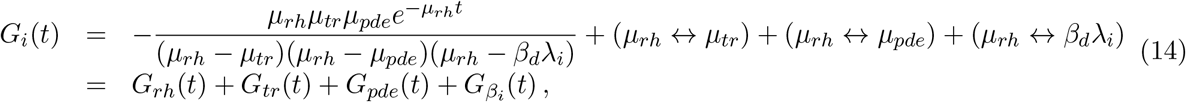

where the double-headed arrows have the same significance as previously. For a step of light at *t* = 0 with duration Δ*t* and constant intensity *ϕ* we get 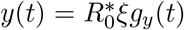 and 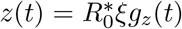, where 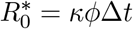 are the number of isomerisations and

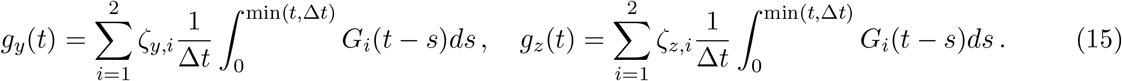

From Eqs. 7-8 we obtain the current 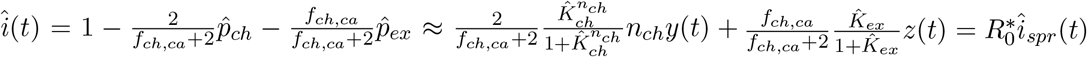, where

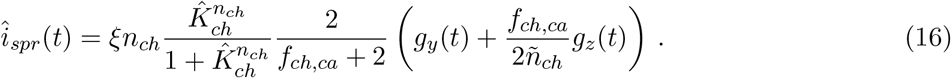

For short flashes with duration 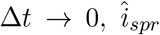 corresponds to the classical single-photon response (SPR). For light steps with duration 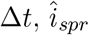 is a unitary response for steps with duration Δ*t*.

We use Eq. 14 to decompose the current into different parts that reflect the contributions from the various biophysical processes,

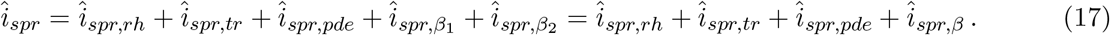

For the SPR with Δ*t* → 0 we have 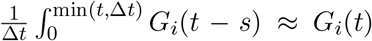, such that 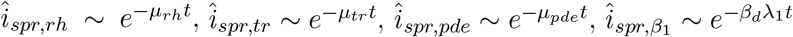, and 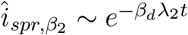.

In the limit *μ*_*ca*_ → ∞ (*r* → ∞) we have 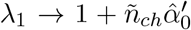 and *λ*_2_ → ∞, and we obtain 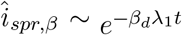, in agreement with [35]. With fast *Ca*^2+^ kinetics we further have 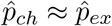 and 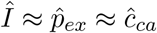 (the latter is valid for 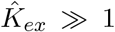), showing that in this limit the *Ca*^2+^ concentration changes in proportion to the current. In the opposite limit *μ*_*ca*_ → 0 (*r* → 0), we have *λ*_1_ → 1, *λ*_2_ → 0, *ζ*_*y*,2_ → 0, *ζ*_*z*,2_ → 0, such that 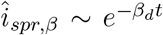. The same behaviour is obtained for the GCAPs^−/−^ case with 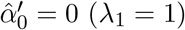.

#### 2.2.1 Emergence of damped oscillations

For 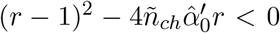, the eigenvalues *λ*_*i*_ are complex conjugate, and 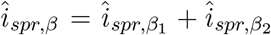 describes a damped oscillation. Such oscillations arise for *r*_1_ *< r < r*_2_, with the bifurcation points 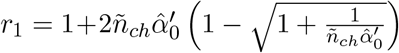 and 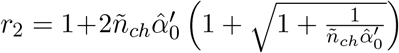. By writing *λ*_1_ = *λ*_*re*_ −*iλ*_*im*_ and *λ*_2_ = *λ*_*re*_ + *iλ*_*im*_, with 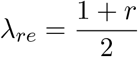 and 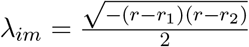, we obtain for a short flash

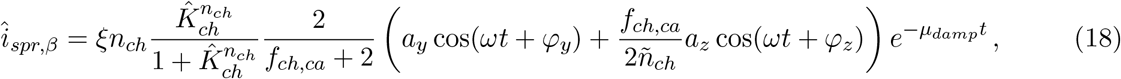

with oscillation frequency 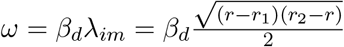 and damping rate 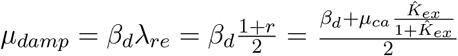. The oscillation amplitudes *a*_*y*_ and *a*_*z*_, and the phases *ϕ*_*y*_ and *ϕ*_*z*_ can be computed with Eq. 14.

#### 2.2.2 Steady state with a dim background light

With constant light intensity *ϕ* we have 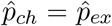 and 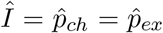. The steady state solutions of Eq. 11 are *z* = *ñ*_*ch*_*y* and 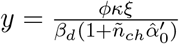. For the current we get 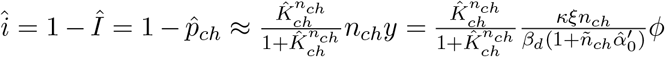.

#### 2.3 Changing the extracellular *Ca*^2+^ concentration

We now investigate how the photoresponse is modified when the extracellular *Ca*^2+^ concentration is altered by a factor of *r*_*ca,ex*_ compared to the previous reference concentration (*r*_*ca,ex*_ = 1). We assume that the CNG current carried by *Ca*^2+^ changes in proportion to the extracellular concentration, 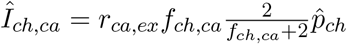, whereas the current not carried by *Ca*^2+^ remains unaffected, 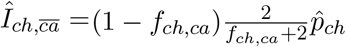. Thus, the CNG current is 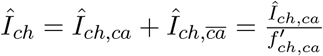, where

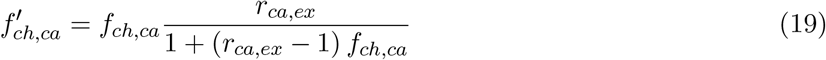

is the new fraction of the CNG current carried by *Ca*^2+^ (*f*_*ch,ca*_ is reference value from Table 1 for *r*_*ca,ex*_ = 1). The overall current is

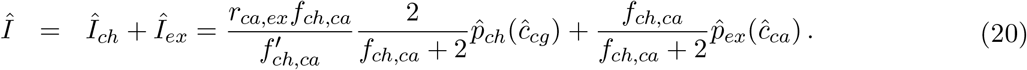

**Table 1:**
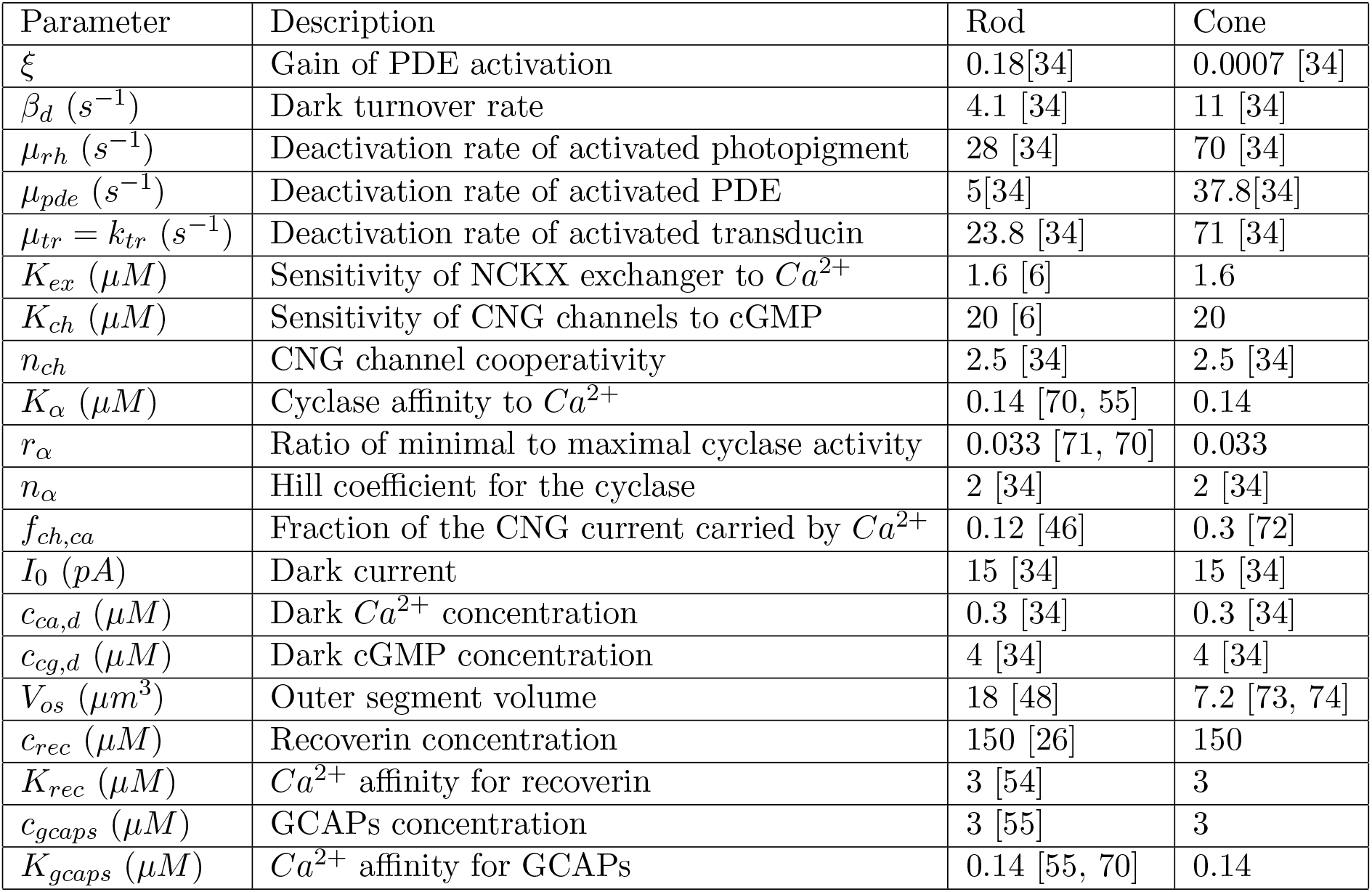
Parameter values for mouse rods and cones. Whenever cone values are not known, we simplify and use the rod values.

With 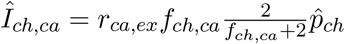, the equation for *ĉ*_*ca*_ now reads

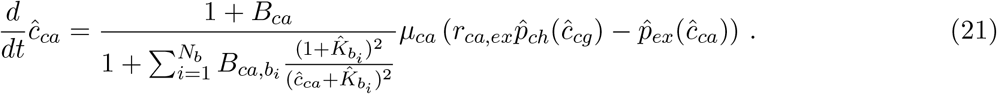

Thus, for *r*_*ca,ex*_ ≠ 1 we have that *ĉ*_*ca*_ = *ĉ*_*cg*_ = 1 are no longer steady state solutions in darkness. The new steady state values *ĉ*_*ca,d*_ and 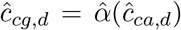 are obtained by solving the equation 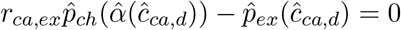. With 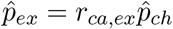 we obtain from Eq. 20 for the dark current

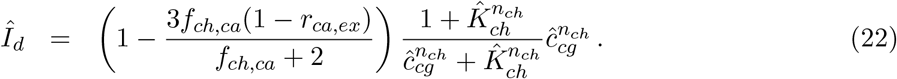

Finally, with 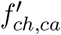, and with parameters that are normalized with the new steady state values *ĉ*_*ca,d*_*c*_*ca*,0_, *ĉ*_*cg,d*_*c*_*cg*,0_ and *Î*_*d*_*I*_0_, we can use Eq. 5 and Eq. 6 with modified parameters to study the response with modified extracellular *Ca*^2+^..

## 3 Results

Eq. 5 reveals that the *Ca*^2+^ kinetics is controlled by the effective rate 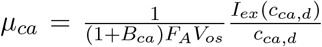 (Eq. 6). The value of *μ*_*ca*_ depends on the effective buffering constant 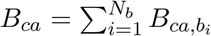, which adds the contributions from the various buffer species. With *B*_*ca*_ = 0 and parameters from Table 1 we compute *μ*_*ca*_ ∼ 1.6 × 10^3^*s*^−1^ for a rod, and *μ*_*ca*_ ∼ 9.4 × 10^3^*s*^−1^ for a cone. With fast *Ca*^2+^ kinetics (large *μ*_*ca*_) we have 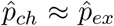, and the time-dependent current can be approximated by the steady state expression 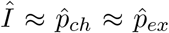. Moreover, for 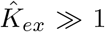 we have 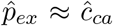 and 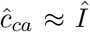. Thus, for large *μ*_*ca*_ and 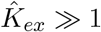 we find that the *Ca*^2+^ concentration changes in proportion to the current. In this limit we recover our results from [34, 35].

We now generalize our previous results by studying the photoresponse as a function of *μ*_*ca*_. A straightforward experimental method to slow down the *Ca*^2+^ dynamics is to add exogenous buffers. This leads to a larger buffering capacity *B*_*ca*_ and a reduced *μ*_*ca*_, but otherwise does not modify Eqs. 5,6. In contrast, modifying exchanger properties affects *μ*_*ca*_ and steady state values. To study such a case, one would have to first adapt Eqs. 5,6. Nevertheless, in the following we use the value of *μ*_*ca*_ to characterize the difference between responses (and not *B*_*ca*_), because it is the value of *μ*_*ca*_ with respect to other rate constants of the model that is relevant.

### 3.1 Validity range of the analytic results

We check the validity range of our asymptotic analysis by comparing simulations and analytic results for 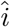. For the simulations, we use Eq. 5 with one low and high affinity buffer each, as specified in Table 1 (recoverin and GCAPs). The buffering parameters in Table 1 are obtained as follows: Recoverin is the most abundant calcium buffer in the outer segment [6, 55], with a dissociation constant *K*_*ca,rec*_ ∼ 3*μM* [54]. To obtain a realistic buffering constant around *B*_*ca*_ ∼ 40 − 50 [47], we estimate that the recoverin concentration has to be around *c*_*rec*_ ∼ 150*μM*. Such values are consistent with estimations from [26, 56]. For GCAPs buffering we use *c*_*gcaps*_ ∼ 3*μM* with a dissociation constant *K*_*ca,gcaps*_ ∼ 0.14*μM* [55]. For the buffering, GCAPs contributes to *B*_*ca*_ only a small value around 3 × 0.14/(0.3 + 0.14)^2^ ∼ 2. Note that we consider the effect of binding between *Ca*^2+^ and recoverin for the buffering, but we neglect the effect of *Ca*^2+^ binding to recoverin for photopigment deactivation. The latter further depends on the interaction between rhodopsin kinase and recoverin, which we assume to occur on a much slower time scale.

To check the validity range of our asymptotic analysis, we compare simulations and analytic results for 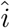 for short flashes and longer step responses with fast and slow *Ca*^2+^ kinetics, characterized by *μ*_*ca*_, for rods (Fig. 1) and cones (Fig. 2). Simulations are computed with Eq. 5, analytic results are 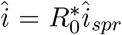, where 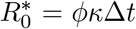 are the number isomerisations produced by *ϕ* during the stimulation time Δ*t*, and 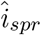 is from Eq. 16. The values for *μ*_*ca*_ displayed in Fig. 1A and Fig. 2B are computed with Eq. 6 using parameters from Table 1. For the simulations with tenfold reduced *μ*_*ca*_, we simply increase both buffer concentrations by a factor of 10.

**Figure 1:**
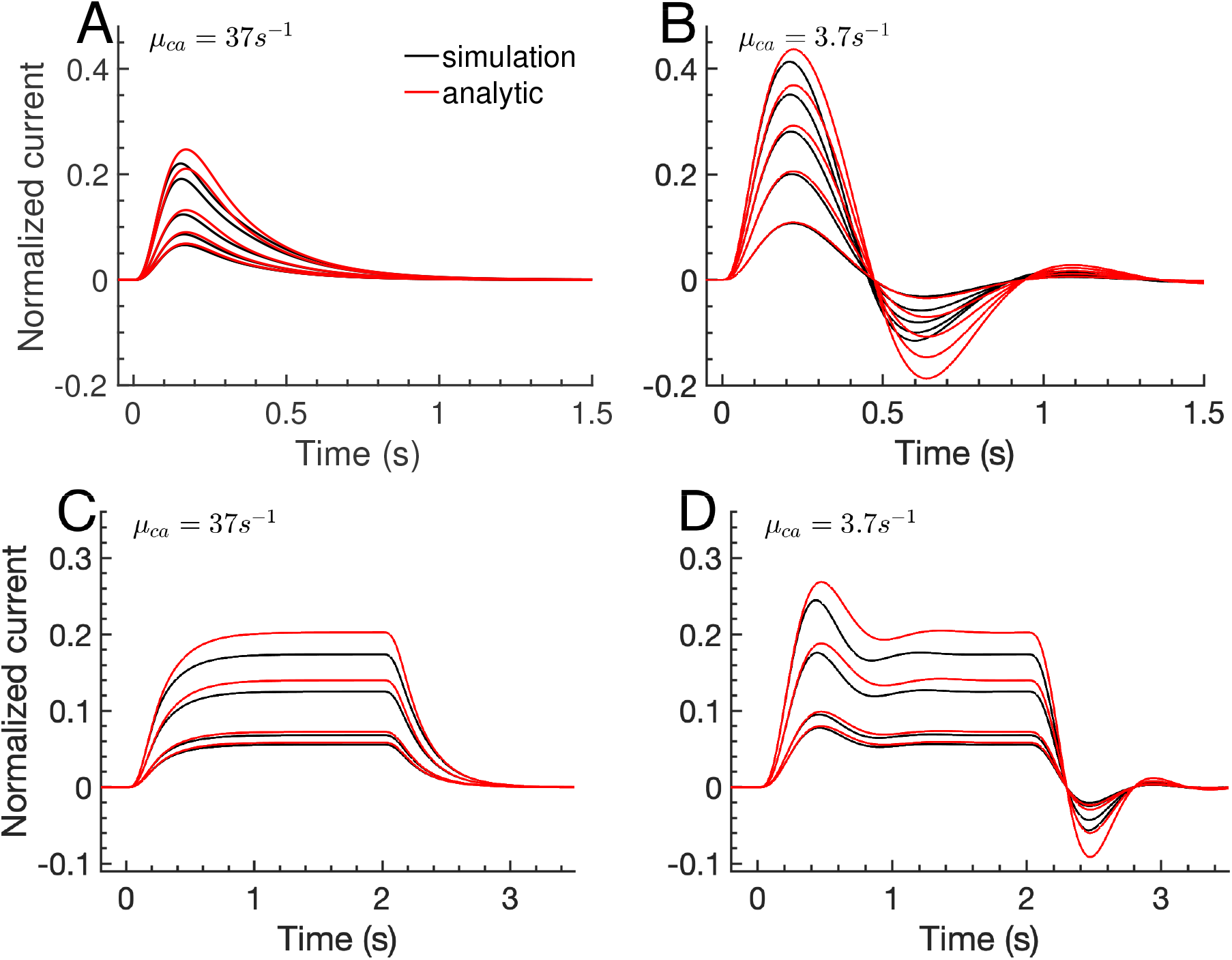
Simulations and analytic results of dim-light responses in a mouse rod. The normalized current 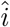 with fast and slow *Ca*^2+^ kinetics is shown for flashes of light at time *t* = 0 with duration Δ*t* = 0.005*s* (A-B), and light steps with duration Δ*t* = 2*s* (C-D). Light intensities are such that the isomerisations 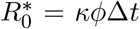 are 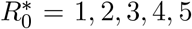 in (A-B), and 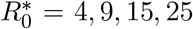 in (C-D). Simulations are performed with Eq. 5 and two *Ca*^2+^ buffers with low and high affinity. In (A,C) the buffer properties are as specified in Table 1 (recoverin and GCAPs buffers). In (B,D) both buffer concentrations have been augmented by a factor of 10. The value of *μ*_*ca*_ shown in the panels is computed with Eq. 6. Analytic results 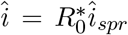 are computed with 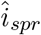 from Eq. 16. If not specified otherwise, parameters are from Table 1.

**Figure 2:**
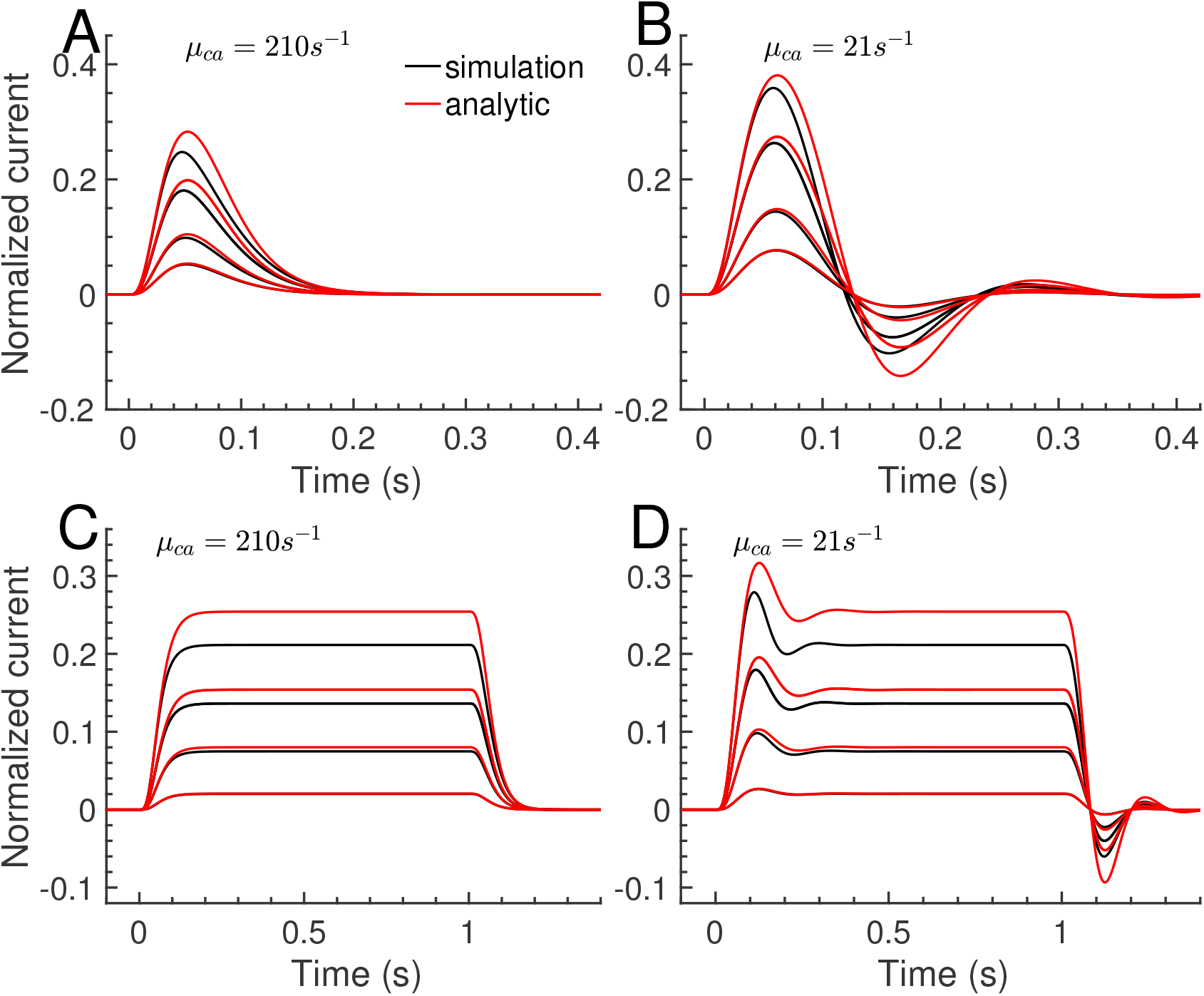
Simulations and analytic results of dim-light responses in a mouse cone. The normalized current 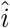 with fast and slow *Ca*^2+^ kinetics is shown for flashes of light at time *t* = 0 with duration Δ*t* = 0.005*s* (A-B), and light steps with duration Δ*t* = 1*s* (C-D). Light intensities are such that the isomerisations 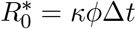 are 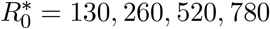 (A-B) and 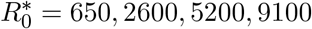 (C-D). Simulations are performed with Eq. 5 and two *Ca*^2+^ buffers with low and high affinity. In (A,C) the buffer concentrations are as specified in Table 1 (recoverin and GCAPs buffers), in (B,D) both buffer concentrations have been augmented by a factor of 10. The value of *μ*_*ca*_ shown in the panels is computed with Eq. 6. Analytic results 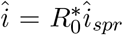 are computed with 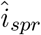 from Eq. 16. If not specified otherwise, parameters are from Table 1.

For dim light, simulations and analytic results for rods and cones faithfully agree for flash responses with fast and slow *Ca*^2+^ dynamics (Fig. 1A-B and Fig. 2A-B, black vs red curves), as well as step responses (Fig. 1C-D and Fig. 2C-D, black vs red curves). As the light intensity increases, the analytic result overestimates the current response because the linear approximation underestimates the non-linear increase in the negative *Ca*^2+^ feedback to the cyclase. For large *μ*_*ca*_ (*μ*_*ca*_ = 37*s*^−1^ for rod, *μ*_*ca*_ = 210*s*^−1^ for cone) we observe monophasic responses (Fig. 1A,C and Fig. 2A,C). With 10-fold increased buffering (*μ*_*ca*_ = 3.7*s*^−1^ for rod, *μ*_*ca*_ = 21*s*^−1^ for cone) we observe damped oscillations (Fig. 1B,D and Fig. 2B,D), as predicted by Eq. 18. In summary, this shows that our analytic formulas can be applied to study the dim-light response.

### 3.2 Single photon response (SPR) and unitary step response

Because dim-light responses are proportional the number of isomerisations 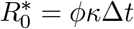, we focus on the unitary response 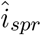 with 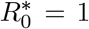 (Eq. 16). For short flashes with Δ*t* → 0, 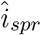 corresponds to the classical single-photon response (SPR). For steps with longer duration Δ*t*, the unitary step response 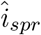 is a response function (a single isomerisation can never produce a step response).

We find that time to peak and amplitude of a SPR decrease as *μ*_*ca*_ increases, because a faster *Ca*^2+^ dynamics leads to quicker cyclase activation and more negative feedback (Fig. 3A,B). Thus, a slower *Ca*^2+^ dynamics implies a larger rod sensitivity (larger amplitude), in agreement with experiments with exogenous buffering [14]. In GCAPs^−/−^ photoreceptors where the *Ca*^2+^ feedback to the cyclase is genetically removed, the *Ca*^2+^ kinetics affects the light response only little via the exchanger current (Fig. 3A-B, dashed lines). This shows that the *Ca*^2+^ dynamics affects the dim-light response mainly because of the cyclase feedback, in agreement with experiments [17].

**Figure 3:**
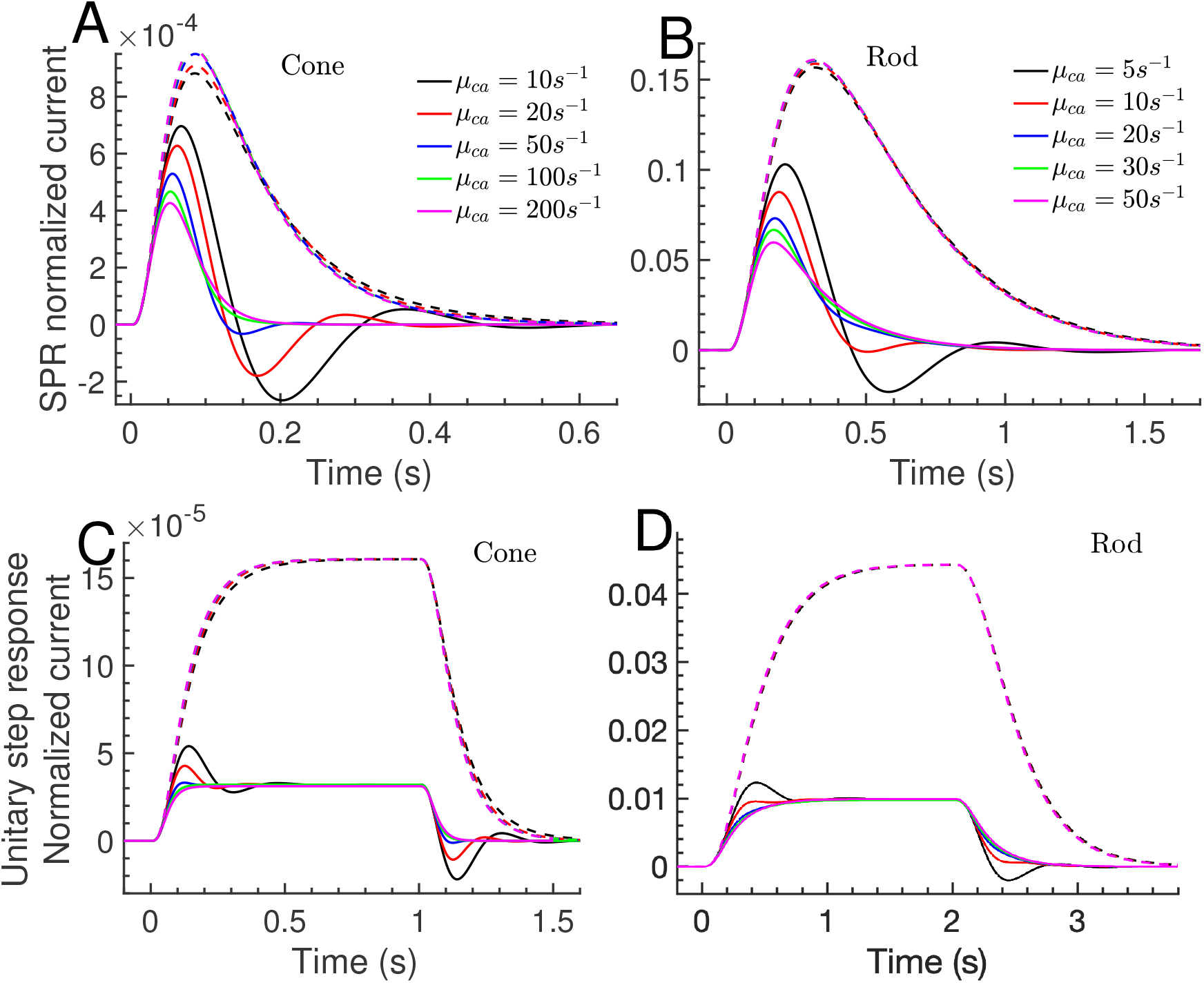
Single-photon responses (SPRs) and unitary step responses for rod and cone. Responses are computed with 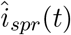 from Eq. 16 and *μ*_*ca*_ values depicted in panel (A) for cone, and in panel (B) for rod. Solid lines show WT, dashed lines GCAPs^−/−^ responses (the *Ca*^2+^ feedback to the cyclase is removed). The cone step responses in (C) have light duration Δ*t* = 1*s*, the rod response in (D) have Δ*t* = 2*s*.

Next we characterize the different phases of a flash and step response. For flash responses, we define the initial phase up to the peak time, and the recovery phase afterwards (Fig. 3A-B). For step responses where Δ*t* is sufficiently long such that an intermediate steady state is reached, we define the initial phase up to the time when the intermediate plateau value is reached; afterwards we define the plateau phase up to the time when the light is switched off, and finally we have the recovery phase (Fig. 3C-D). Slowing down the *Ca*^2+^ kinetics increases both amplitude and time to peak of a SPR, and damped oscillations start to appear during the recovery phase (Fig. 3A-B) (see the analysis in section 2.2.1). For a step response, only initial and recovery phases depend on *μ*_*ca*_, whereas the plateau phase is determined by steady-state properties that do not depend on *μ*_*ca*_ (Fig. 3C-D, solid lines). We compute that the GCAPs^−/−^ plateau current is by a factor of 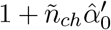 larger compared to the WT value (see the analysis after Eq. 18). With Table 1, we find 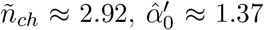 and 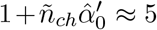. Thus, whereas the plateau currents between WT and GCAPs^−/−^ step responses differ by a factor of around 5 (Fig. 3C-D), the WT and GCAPs^−/−^ SPR amplitudes depend on *μ*_*ca*_ and differ maximally by a factor of around 2-3 with fast *Ca*^2+^ kinetics (Fig. 3A-B).

### 3.3 Waveform of flash and step responses

Changing the *Ca*^2+^ kinetics affects amplitude and shape of flash and step responses (Fig. 3). To compare how the response shape is affected, we use waveforms where the response amplitude is normalized to one. For short flashes (see Fig. 3A-B), we define the current waveform by normalizing 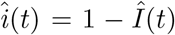 with the peak amplitude. For longer step responses (see Fig. 3C-D), we define the waveform by normalizing 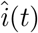 with the intermediate plateau value. To compare current and *Ca*^2+^ changes, we define the *Ca*^2+^ waveform using the change in the *Ca*^2+^ concentration 1 − *ĉ*_*ca*_(*t*). For dim light we have 1 − *ĉ*_*ca*_(*t*) ≈ *z*(*t*).

The current waveform is monophasic for large *μ*_*ca*_ (Fig. 4A-B, magenta curves), and in this limit the *Ca*^2+^ concentration changes in proportion to the current (Fig. 4C-D, green curves). As *μ*_*ca*_ decreases, damped oscillations emerge (Fig. 4A-B) because the *Ca*^2+^ dynamics becomes more and more delayed with respect to the current (Fig. 4C-D, red and black curves). In limit *μ*_*ca*_ → 0 the current response becomes monophasic because *Ca*^2+^ concentration and cyclase activity remain constant (nor shown, but see also the GCAPs^−/−^ responses in Fig. 3).

**Figure 4:**
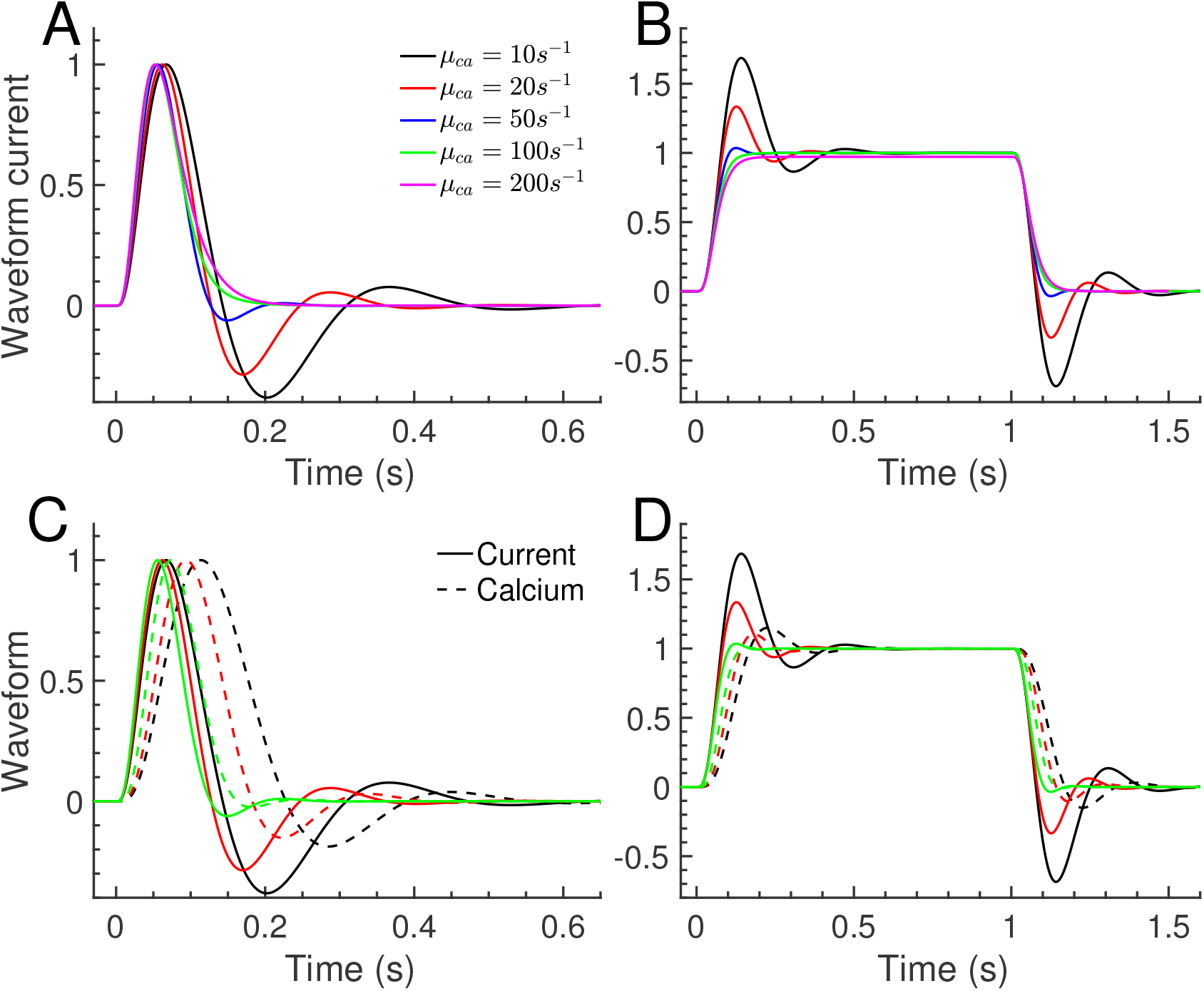
Waveform of flash and step responses in cone. (A) The current waveform of a dim-flash response is obtained by normalizing the SPRs from Fig. 3A with the peak amplitude. (B) For longer step responses, the waveform is obtained by normalizing the responses in Fig. 3C with the intermediate plateau values. (C-D) The waveform of the *Ca*^2+^ concentration is computed similar to the current waveform using 1 − *ĉ*_*a*_(*t*). The current waveforms (solid lines) from (A-B) for *μ*_*ca*_ = 10, 20, 50*s*^−1^ are compared to the corresponding *Ca*^2+^ waveforms (dashed lines). The color coding is as in (A-B).

Whereas the flash shape differs between initial and recovery phase (Fig. 4A), there is a symmetry between these phases for step responses (Fig. 4B). Indeed, when an intermediate steady state is reached, we find from Eq. 11 that initial and recovery phase are related by 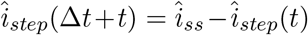, where 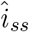 is the intermediate steady state current.

### 3.4 Waveform decomposition

Next we use the analytic result in Eq. 17 to show how the various biophysical processes contribute to the response. Such information cannot be obtained from simulations. We find that the recovery in a rod with fast *Ca*^2+^ dynamics is limited by PDE decay, 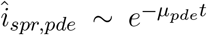 (Fig. 5A)). For a GCAPs^−/−^ cone we have that the recovery is limited by the basal turnover rate 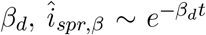 (Fig. 5B. For these two cases, parameters can be extracted by fitting the response recovery with a single exponential. In general, however, the recovery of flash responses is not determined by a single exponential. For example, with slow *Ca*^2+^ dynamics the recovery phase contains a damped oscillations (Fig. 5C-D), in which case fitting the response decay with a single exponential does not yield meaningful parameter values.

**Figure 5:**
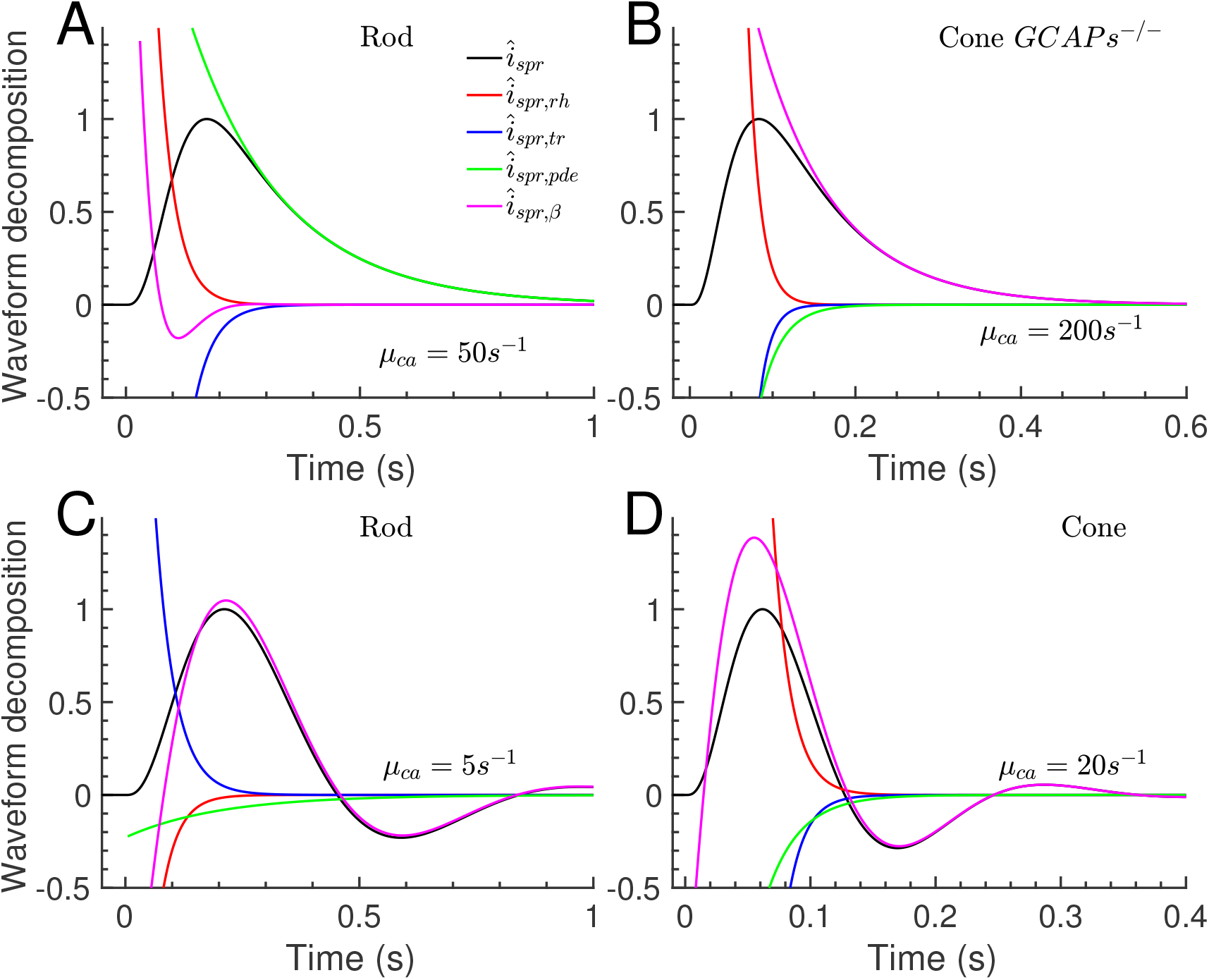
Decomposition of the SPR waveform. The decomposition of the SPR waveform (black line) is computed with Eq. 17. In every panel, the black line is the sum of the colored lines. (A) SPR decomposition for a WT rod with *μ*_*ca*_ = 50*s*^−1^, (B) GCAPs^−/−^ cone with *μ*_*ca*_ = 20*s*^−1^, (C) WT rod with *μ*_*ca*_ = 5*s*^−1^, (D) WT cone with *μ*_*ca*_ = 20*s*^−1^.

### 3.5 Damped oscillations

The analysis in section 2.2.1 shows that the response contains a damped oscillation if the ratio 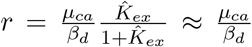 is within the range *r*_1_ *< r < r*_2_, where 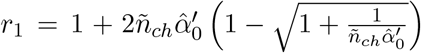 and 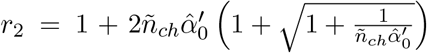 depend on the CNG channel cooperativity *n*_*ch*_ and the sensitivity of the cyclase to 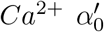 (see section 2.2.1). For example, with Table 1 we compute 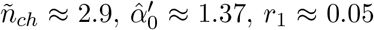 and *r*_2_ ≈ 18. The rage for *μ*_*ca*_ where oscillations occur, determined by 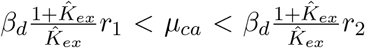, increases as a function of the channel cooperativity and cyclase feedback 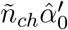 (Fig. 6A; cone (black curve) and rod (red curve)). This range is wider for a cone because of the larger *β*_*d*_ (Fig. 6A). No oscillations occur for Gcaps^−/−^ mutants with 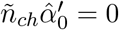. The oscillation frequency depends on 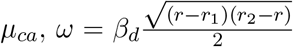 (Eq. 18). As function of *μ*_*ca*_, *ω* increases from zero towards the maximal frequency 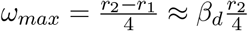, which is attained for 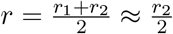 and 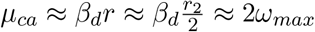. Afterwards, *ω* decreases again as *μ*_*ca*_ further increases (Fig. 6B).

**Figure 6:**
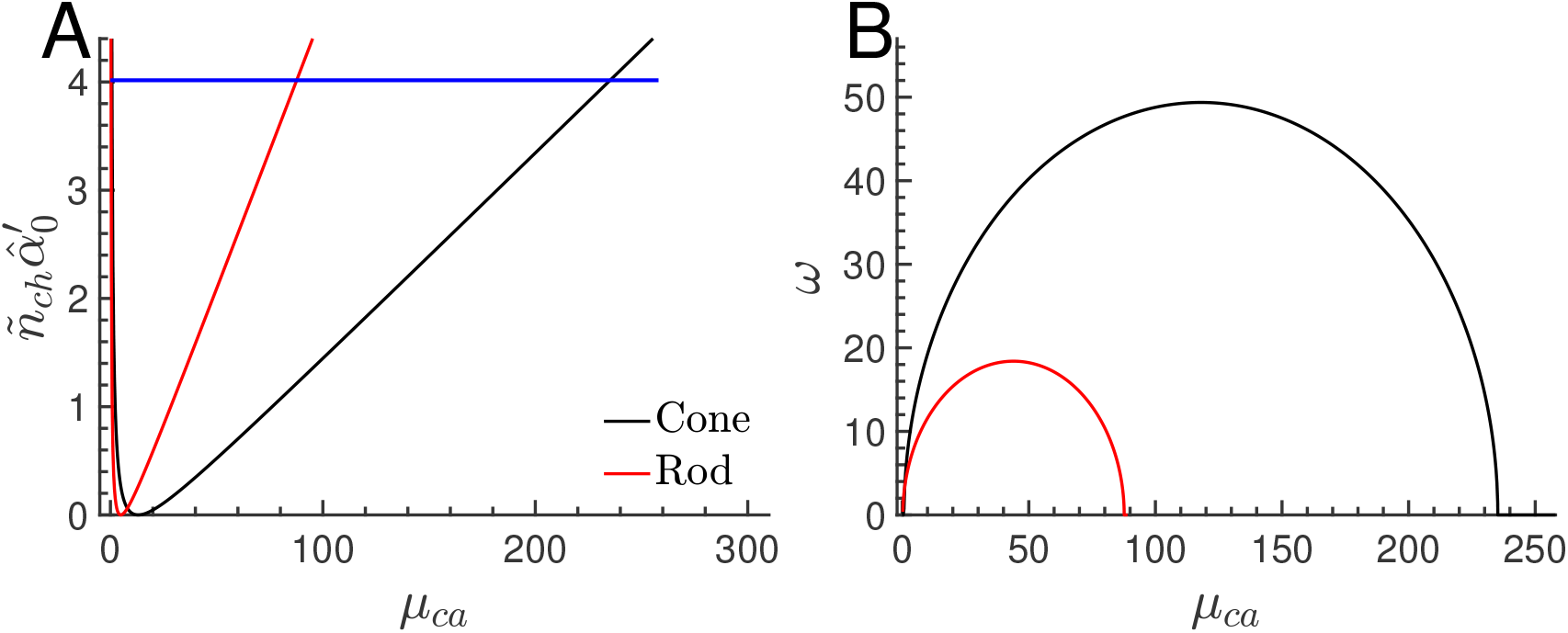
Phase space for *Ca*^2+^ oscillations. (A) Range for *μ*_*ca*_ where oscillations occur as a function of 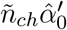 for rod (red) and cone (back) (see section 2.2.1 for more details). The blue horizontal line indicates the value 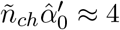 computed with parameters from Table 1. (B) Oscillation frequency *ω* as a function of *μ*_*ca*_ for rod and cone. The maximal frequency 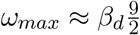 is achieved for *μ*_*ca*_ ≈ *β*_*d*_9.

Although oscillations occur over a wide range of *μ*_*ca*_ values (Fig. 6A), they affect the recovery phase only if the damping rate 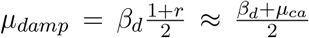 is not too large. Indeed, for large *μ*_*ca*_ oscillations are damped before they reach the recovery phase (Fig. 5A, magenta curve). As *μ*_*ca*_ decreases, oscillations start to affect the recovery phase (Fig. 5C-D). It is difficult to give a precise value for *μ*_*ca*_ that indicates when oscillations start to become visible during the recovery phase, which depends on damping but also the initial amplitude of the oscillation. We note, however, that for large 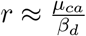 where the *Ca*^2+^ dynamics is determined by the current, the response changes only slowly as a function of *r* (see Fig. 3A-B). More profound changes start to occur for 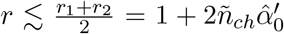 where 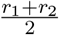 corresponds to the center of the oscillation ranges Fig. 6A. From this we obtain that the response becomes significantly altered for *μ*_*ca*_ ≲ *β*_*d*_*r*^*∗*^ ≈ 9*β*_*d*_, which seems compatible with Fig. 3A-B.

### 3.6 Effect of changing the extracellular *Ca*^2+^ concentration

Finally, we explored how changing the extracellular *Ca*^2+^ concentration affects the light response in a cone (similar conclusions apply for a rod, not shown). We consider that the extracellular concentration is altered by a factor of *r*_*ca,ex*_ compared to the previous reference value, which alters the fraction of the CNG current that is carried by *Ca*^2+^ by a factor of 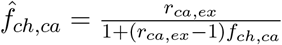 (see Eq. 19), where *f*_*ch,ca*_ = 0.3 is the reference value for *r*_*ca,ex*_ = 1. As a consequence of the modified *Ca*^2+^ influx, the dark steady-state values *ĉ*_*ca,d*_, *ĉ*_*cg,d*_ and *Î*_*d*_ also change (Fig. 7A, black, red and blue curves). For example, when the extracellular *Ca*^2+^ concentration is lowered tenfold (*r*_*ca,ex*_ = 0.1), the fraction of the CNG current that is carried by *Ca*^2+^ is reduced by a factor of around 7 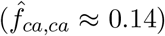. The dark *Ca*^2+^ concentration in the OS is reduced by a factor of around 2 (*ĉ*_*ca,d*_ ≈ 0.55), which activates the cyclase and results in an around 2.1-fold higher cGMP concentration (*ĉ*_*cg,d*_ ≈ 2.1), and an around 3.8-fold larger dark current (*Î*_*d*_ ≈ 3.8). In contrast, the value of *μ*_*ca*_ changes only little because 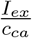 is almost constant for *K*_*ex*_ ≫ *c*_*ca*_, and because recoverin has a low *Ca*^2+^ affinity (Table 1), such that *B*_*ca*_ is only little affected by the reduction in *Ca*^2+^.

**Figure 7:**
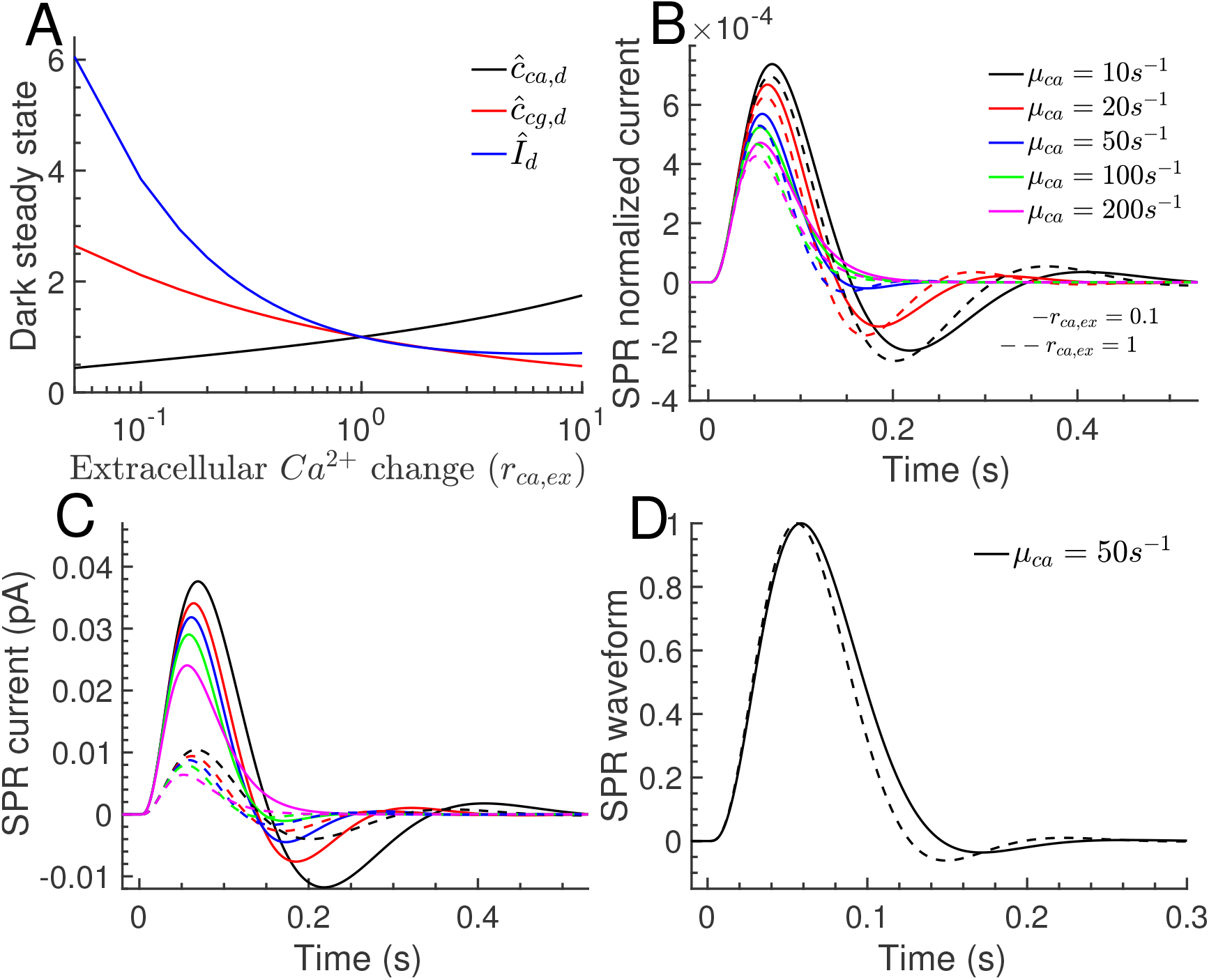
Effect of changing the extracellular *Ca*^2+^ concentration for a cone. (A) Changes in the dark steady state concentrations of *Ca*^2+^ (black curve), cGMP (red curve), and dark current (blue curve) when the extracellular *Ca*^2+^ concentration is altered by a factor of *r*_*ca,ex*_. (B) Comparison of the normalized SPR current 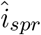 (Eq. 16) for *r*_*ca,ex*_ = 0.1 (solid lines) and *r*_*ca,ex*_ = 1 (dashed lines) for various *μ*_*ca*_. The dashed curves are identical to the solid curves in Fig. 3A. (C) The dimensional SPR currents are obtained by multiplying the traces in (B) with the corresponding dark currents *I*_*d*_ = 3.8 × 15*pA* (*r*_*ca,ex*_ = 0.1) and *I*_*d*_ = 15*pA* (*r*_*ca,ex*_ = 1). (D) Comparison of the SPR waveforms for *μ*_*ca*_ = 50*s*^−1^.

To study the dynamics of 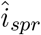 with *r*_*ca,ex*_ = 0.1, we use Eq. 16 with parameters that are normalized with the new steady state values, and with the new value for *f*_*ch,ca*_. We find that a tenfold reduction of extracellular *Ca*^2+^ slightly increases the normalized amplitude and the duration of the SPR (Fig. 7B; solid curves are for *r*_*ca,ex*_ = 0.1, dashed curves for *r*_*ca,ex*_ = 1). Since *μ*_*ca*_ is not much changed, this effect is due to the smaller cyclase feedback 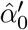, which is reduced from the reference value 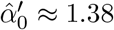 (*r*_*ca,ex*_ = 1) to 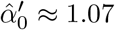 (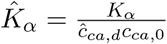 is larger for *r*_*ca,ex*_ = 0.1). Although the normalized responses are not much different, because of the 3.8-fold larger dark current (*I*_*d*_ = 15*pA* for *r*_*ca,ex*_ = 1 (Table 1); *I*_*d*_ = 3.8 × 15*pA* for *r*_*ca,ex*_ = 0.1) the amplitude of the SPR current 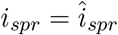 is much increased with reduced extracellular *Ca*^2+^ (Fig. 7C, solid vs dashed curves). By comparing the current waveforms for *μ*_*ca*_ = 50*s*^−1^, we find the the undershoot is slightly reduced for *r*_*ca,ex*_ = 0.1 (Fig. 7D).

## 4 Discussion

The signal transduction pathway of rod and cone photoreceptors consists of a cascade of biophysical processes that transform light into a current response. Many of these steps depend on *Ca*^2+^ feedback, which can be manipulated by exogenous buffers, or by changing the extracellular *Ca*^2+^ concentration.

So far, the impact of *Ca*^2+^ on the response dynamics has mostly been studied using numerical simulations [36, 37, 57, 44, 58, 27]. In this work, we derived analytic results for dim-light responses in rods and cones that show how response sensitivity and dynamics change as the *Ca*^2+^ kinetics is altered due to fast buffering and change in the extracellular *Ca*^2+^ concentration. Our analytic formulas not only provide quantitative and conceptual insight, but they are also a means to easily compute photoreceptor responses based on underlying biophysical parameters without having to numerically solve a system of differential equations, which can be used as input to study downstream processes in the retina [59].

We found that the *Ca*^2+^ kinetics in presence of fast buffering is controlled the effective rate 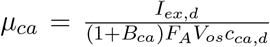 (Eq. 6). *B*_*ca*_ is an effective buffering constant that has contributions from the various buffer species (Eq. 6), *V*_*os*_ is the outer segment volume, *I*_*ex,d*_ is the exchanger current in darkness, *c*_*ca,d*_ is the dark *Ca*^2+^ concentration, and *F*_*A*_ is the Faraday constant. For large *μ*_*ca*_, light responses are monophasic (Fig. 3, magenta curves) and the *Ca*^2+^ concentration changes in proportion to the current (Fig. 4C-D, green curves). As *μ*_*ca*_ decreases, the *Ca*^2+^ concentration becomes delayed with respect to the current (Fig. 4C-D, red and black curves), and damped oscillations emerge (Fig. 4A-B). Our analysis shows that fast buffering alone can induce oscillations, and the presence of slow buffers is not mandatory.

The phase space analysis reveals that the transition form monophasic to biphasic (oscillatory) responses depends on the ratio *μ*_*ca*_*/β*_*d*_. Oscillations occur if this ratio is below a threshold value of around 18, which depends on the strength of the cyclase feedback 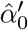, and the channel cooperativity *n*_*ch*_. Because *β*_*d*_ is much larger in a mouse cone compared to rod [34], the range for *μ*_*ca*_ where oscillations can occur is much larger in a cone (Fig. 6A). The differences in *β*_*d*_ might be one of the reasons why oscillations are rarely observed in amphibian compared to primate rods [37], and why they have been more frequently observed in cones [29, 30, 31]. We further computed that oscillations are damped with rate 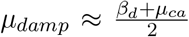, and we determined the oscillation frequency as a function of *μ*_*ca*_ (Fig. 6B). Although the range of *μ*_*ca*_ where oscillations occur is large (Fig. 6A), oscillations are visible during the recovery phase of flash responses only if the damping is not too strong, which requires that the *Ca*^2+^ kinetics is sufficiently slowed down (compare the magenta curves in Fig. 5A,C). With the analytic results it is possible to decompose the dim-flash response into individual terms that reflect contributions from various biophysical processes (Eq. 17 and Fig. 5). Such a decomposition cannot be performed with simulations only. As expected, the decomposition shows that oscillations are present in the term that describes the interaction between *Ca*^2+^ and cGMP synthesis (Fig. 5, magenta curves). In GCAPs^−/−^ photoreceptors where this interaction is absent, flash responses are monophasic, and they vary only little with *μ*_*ca*_ (Fig. 3 dashed lines), in agreement with experiments [17]. The small dependency on *μ*_*ca*_ for GCAPs^−/−^ photoreceptors is a consequence of *Ca*^2+^-dependent exchanger current. In WT photoreceptors with intact cyclase feedback, modifying the *Ca*^2+^ kinetics strongly affects amplitude and shape of the light response (Fig. 3, solid lines). The sensitivity (amplitude) of the single-photon response (SPR) increases gradually as the *Ca*^2+^ kinetics is slowed down (Fig. 3A-B, solid lines), which is because the negative *Ca*^2+^ feedback to the cyclase becomes delayed (Fig. 4C). This agrees with larger photoreceptor sensitivities observed with exogenous buffering [14]. We find a maximal sensitivity ratio around 2-3 between GCAPs^−/−^ and WT photoreceptors for fast *Ca*^2+^ kinetics. Because the observed sensitivity ratio between GCAPs^−/−^ and WT photoreceptors is around 3 [34, 17, 52], this suggests that under physiological conditions the *Ca*^2+^ kinetics is fast, and that the *Ca*^2+^ concentration changes in proportion to the current [51, 60]. For step responses with an intermediate plateau phase, oscillations are visible during the initial and recovery phase of the response (Fig. 4B). The plateau value is determined by steady-state properties and independent of the *Ca*^2+^ kinetics. We computed that the plateau value in a GCAPs^−/−^ photoreceptor is by a factor of 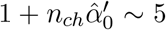 larger compared to WT (Fig. 3C-D). Since this ratio is independent of the *Ca*^2+^ kinetics, it could be used to estimate the strength of the cyclase feedback 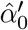. Whereas initial and recovery phase of flash responses are different (Fig. 4A), there is a symmetry between these phases for step responses (Fig. 4B); see also Fig. 5B in [61]). Hence, the analysis of initial and recovery phase of dim step responses provides the same information, whereas complementary information can be extracted from flash response (see [35]).

We explored how reducing the extracellular *Ca*^2+^ concentration affects the SPR response in a cone (similar conclusions apply to rod, not shown). Modifying extracellular *Ca*^2+^ alters the fraction *f*_*ch,ca*_ of the CNG current that is carried by *Ca*^2+^ (Eq. 19), and thereby changes the steady state values of *Ca*^2+^, cGMP and current (Fig. 7A). For example, a tenfold reduction in extracellular *Ca*^2+^ reduces the dark *Ca*^2+^ concentration by a factor of 2, it increases the SPR amplitude by a factor of (Fig. 7A), and it prolongs the SPR duration (Fig. 7B). Similar changes have been observed experimentally for primate cones when the extracellular *Ca*^2+^ concentration has been reduced (Fig. 2A in [33]). In [33] it has also occasionally been observed that a biphasic response could be transformed into a monophasic response by lowering the extracellular *Ca*^2+^ concentration (Fig. 2A in [33]). Although we also observe a small reduction of the undershoot (Fig. 7C), the effect seems much weaker than shown in [33]. A possible reason for this could be that, occasionally, the reduction in the extracellular *Ca*^2+^ concentration leads to a complete saturation of the cyclase, which abolishes the oscillation and gives monophasic responses.

In this work we did not consider slow buffers in order to keep the analysis most comprehensible. However, it is straightforward to generalize the analysis with slow buffering in future work. In first approximation, and with a single slow buffer species, Eq. 11 will be replaced by 3-dimensional system of equations. Although the phase space is now more complex, we do not expect to find solutions that are qualitatively much different from what we have observed here, which is consistent with simulations in presence of slow buffering reactions [36, 37]

We used a non-spatial (well-stirred) model for the outer segment, which allowed to derive analytic results. Non-spatial models are frequently used to study the photoresponse [34, 62, 42, 41, 27, 40, 63, 58, 64, 44, 43, 36]. Whereas spatial models are a more accurate description of reality, the downside is that no analytic results can usually be derived [65, 49, 48, 46, 66, 67]. Nevertheless, we performed simulations with our spatial model from [48] to check that our conclusions remain valid (not shown). Recently it has been found that the large outer segment of a frog rod is a spatially inhomogeneous compartment [60, 68]. It remains unclear whether this is also true for the much smaller outer segment of mouse or primate rods and cones. We leave it for future work to investigate how a possible spatial inhomogeneity might affect the photoresponse.

Although we focused on the dim-light response in darkness, the linear response analysis that we presented can be easily generalized to the case with a background light. In this case, one has to first numerically compute the steady-state values corresponding to the background light, and then use the analysis with parameters that have been normalized with these steady state values.

In this work we performed mathematical analysis to obtain quantitative and conceptual understanding. We hope that this work will serve as a template to derive more general results in future work, and that it might inspire new experiments to test the *Ca*^2+^ dynamics. Apart from phototransduction, our analysis might also be applied to study other G-protein coupled signalling cascades, for example, in olfaction [69].

## Author contributions

AA performed analysis and generated figures. JR designed the research, performed analysis and wrote the manuscript.

## Acknowledgements

We are grateful to Gordon Fain for comments and careful reading of the manuscript. This work was funded by a grant from Agence Nationale de Recherche (ANR) to JR (ANR-19-CE45-0004-01).

## References

[1] T. Ebrey and Y. Koutalos, “Vertebrate photoreceptors,” Progress in retinal and eye research, vol. 20, no. 1, pp. 49–94, 2001.

[2] M. Burns and D. Baylor, “Activation, deactivation, and adaptation in vertebrate photoreceptor cells,” Annu. Rev. Neurosci., vol. 24, pp. 779–805, 2001.

[3] M. E. Burns and E. N. Pugh Jr, “Lessons from photoreceptors: turning off g-protein signaling in living cells,” Physiology, vol. 25, no. 2, pp. 72–84, 2010.

[4] V. Arshavsky and M. Burns, “Photoreceptor signaling: supporting vision across a wide range of light intensities.,” J Biol Chem, vol. 287, pp. 1620–1626, 2012.

[5] V. Arshavsky, T. Lamb, and E. Pugh Jr, “G proteins and phototransduction,” Annu. Rev. Physiol., vol. 64, pp. 153–187, 2002.

[6] E. Pugh Jr and T. Lamb, “Phototransduction in vertebrate rods and cones: Molecular mechanism of amplification, recovery and light adaptation,” In Handbook of Biological Physics Vol. 3, Elsevier Science B. V., Amsterdam, pp. 183–255, 2000.

[7] F. Vinberg, J. Chen, and V. Kefalov, “Regulation of calcium homeostasis in the outer segments of rod and cone photoreceptors.,” Prog Retin Eye Res., vol. 67, pp. 87–101, 2018.

[8] K.-W. Koch and D. DellOrco, “A calcium-relay mechanism in vertebrate phototransduction,” ACS chemical neuroscience, vol. 4, no. 6, pp. 909–917, 2013.

[9] K. Nakatani, C. Chen, K. Yau, and Y. Koutalos, “Calcium and phototransduction.,” Adv Exp Med Biol., vol. 514, pp. 1–20, 2002.

[10] J. Korenbrot and T. Rebrik, “Tuning outer segment ca2+ homeostasis to phototransduction in rods and cones.,” Adv Exp Med Biol., vol. 514, pp. 179–203, 2002.

[11] G. Rispoli, “Calcium regulation of phototransduction in vertebrate rod outer segments,” Journal of Photochemistry and Photobiology B: Biology, vol. 44, no. 1, pp. 1–20, 1998.

[12] H. Matthews, R. Murphy, G. Fain, and T. Lamb, “Photoreceptor light adaptation is mediated by cytoplasmic calcium concentration,” Nature, vol. 334, no. 6177, pp. 67–69, 1988.

[13] K. Nakatani and K.-W. Yau, “Calcium and light adaptation in retinal rods and cones,” Nature, vol. 334, no. 6177, pp. 69–71, 1988.

[14] H. Matthews, “Incorporation of chelator into guinea-pig rods shows that calcium mediates mammalian photoreceptor light adaptation.,” The Journal of physiology, vol. 436, no. 1, pp. 93–105, 1991.

[15] G. Fain, H. Matthews, M. Cornwall, and Y. Koutalos, “Adaptation in vertebrate photoreceptors,” Physiol. Reviews, vol. 81, no. 1, pp. 117–151, 2001.

[16] E. N. Pugh Jr, S. Nikonov, and T. Lamb, “Molecular mechanisms of vertebrate photoreceptor light adaptation,” Current opinion in neurobiology, vol. 9, no. 4, pp. 410–418, 1999.

[17] M. E. Burns, A. Mendez, J. Chen, and D. A. Baylor, “Dynamics of cyclic gmp synthesis in retinal rods.,” Neuron., vol. 36, pp. 81–91, 2002.

[18] K. Sakurai, J. Chen, and V. Kefalov, “Role of guanylyl cyclase modulation in mouse cone phototransduction.,” J. Neurosci., vol. 31, pp. 7991–8000, 2011.

[19] Y. Koutalos, K. Nakatani, and K. Yau, “The cgmp-phosphodiesterase and its contribution to sensitivity regulation in retinal rods.,” The Journal of general physiology, vol. 106, no. 5, pp. 891– 921, 1995.

[20] V. Torre, H. Matthews, and T. Lamb, “Role of calcium in regulating the cyclic gmp cascade of phototransduction in retinal rods.,” Proceedings of the National Academy of Sciences, vol. 83, no. 18, pp. 7109–7113, 1986.

[21] V. Torre, H. Matthews, and T. Lamb, “Role of calcium in regulating the cyclic gmp cascade of phototransduction in retinal rods,” Proceedings of the National Academy of Sciences, vol. 83, no. 18, pp. 7109–7113, 1986.

[22] T. Lamb, H. Matthews, and V. Torre, “Incorporation of calcium buffers into salamander retinal rods: a rejection of the calcium hypothesis of phototransduction.,” The Journal of Physiology, vol. 372, no. 1, pp. 315–349, 1986.

[23] J. I. Korenbrot and D. L. Miller, “Cytoplasmic free calcium concentration in dark-adapted retinal rod outer segments,” Vision research, vol. 29, no. 8, pp. 939–948, 1989.

[24] F. Rieke and D. Baylor, “Origin of reproducibility in the responses of retinal rods to single photons,” Biophys. J., vol. 75, pp. 1836–1857, 1998.

[25] G. Field and F. Rieke, “Mechanisms regulating variability of the single-photon responses of mammalian rod photoreceptors,” Neuron, vol. 35, pp. 733–747, 2002.

[26] C. L. Makino, R. Dodd, J. Chen, M. E. Burns, A. Roca, M. Simon, and D. Baylor, “Recoverin regulates light-dependent phosphodiesterase activity in retinal rods,” The Journal of general physiology, vol. 123, no. 6, pp. 729–741, 2004.

[27] J. Korenbrot, “Speed, adaptation, and stability of the response to light in cone photoreceptors: the functional role of ca-dependent modulation of ligand sensitivity in cgmp-gated ion channels.,” J Gen Physiol, vol. 139, no. 1, pp. 31–56, 2012.

[28] D. Holcman and J. Korenbrot, “The limit of photoreceptor sensitivity: Molecular mechanism of dark noise in retinal cones,” J. Gen. Physiol., vol. 125, pp. 641–660, 2005.

[29] D. Schneeweis and J. Schnapf, “The photovoltage of macaque cone photoreceptors: adaptation, noise, and kinetics.,” J Neurosci., vol. 19, pp. 1203–16, 1999.

[30] J. Schnapf, B. Nunn, M. Meister, and D. Baylor, “Visual transduction in cones of the monkey macaca fascicularis,” J. Physiol., vol. 427, pp. 681–713, 1990.

[31] D. Baylor, B. Nunn, and J. Schnapf, “Spectral sensitivity of cones of the monkey macaca fascicularis.,” The Journal of Physiology, vol. 390, no. 1, pp. 145–160, 1987.

[32] H. Matthews, G. Fain, R. Murphy, and T. Lamb, “Light adaptation in cone photoreceptors of the salamander: a role for cytoplasmic calcium.,” The Journal of Physiology, vol. 420, no. 1, pp. 447–469, 1990.

[33] L.-H. Cao, D.-G. Luo, and K.-W. Yau, “Light responses of primate and other mammalian cones,” Proceedings of the National Academy of Sciences, vol. 111, no. 7, pp. 2752–2757, 2014.

[34] J. Reingruber, N. Ingram, K. Griffis, and G. Fain, “A kinetic analysis of mouse rod and cone photoreceptor responses.,” The Journal of Physiology, vol. 598, no. 17, pp. 3747–63, 2020.

[35] A. Abtout, G. Fain, and J. Reingruber, “Analysis of waveform and amplitude of mouse rod and cone flash responses.,” The Journal of Physiology, vol. 599, no. 13, pp. 3295–3312, 2021.

[36] S. Forti, A. Menini, G. Rispoli, and V. Torre, “Kinetics of phototransduction in retinal rods of the newt triturus cristatus.,” The Journal of Physiology, vol. 419, no. 1, pp. 265–295, 1989.

[37] T. Tamura, K. Nakatani, and K. W. Yau, “Calcium feedback and sensitivity regulation in primate rods.,” The Journal of general physiology, vol. 98, no. 1, pp. 95–130, 1991.

[38] R. Hamer, S. Nicholas, D. Tranchina, P. Liebman, and T. Lamb, “Multiple steps of phosphorylation of activated rhodopsin can account for the reproducibility of vertebrate rod single-photon responses,” J. Gen. Physiol., vol. 122, pp. 419–444, 2003.

[39] D. Dell?Orco, H. Schmidt, S. Mariani, and F. Fanelli, “Network-level analysis of light adaptation in rod cells under normal and altered conditions,” Molecular bioSystems, vol. 5, no. 10, pp. 1232– 1246, 2009.

[40] J. Chen, M. Woodruff, T. Wang, F. Concepcion, D. Tranchina, and G. Fain, “Channel modulation and the mechanism of light adaptation in mouse rods.,” J Neurosci, vol. 30, no. 48, pp. 16232–16240, 2010.

[41] B. M. Invergo, D. Dell’Orco, L. Montanucci, K.-W. Koch, and J. Bertranpetit, “A comprehensive model of the phototransduction cascade in mouse rod cells,” Molecular BioSystems, vol. 10, no. 6, pp. 1481–1489, 2014.

[42] L. Astakhova, M. Firsov, and V. Govardovskii, “Activation and quenching of the phototransduction cascade in retinal cones as inferred from electrophysiology and mathematical modeling,” Molecular vision, vol. 21, p. 244, 2015.

[43] J. Sneyd and D. Tranchina, “Phototransduction in cones: an inverse problem in enzyme kinetics.,” Bull Math Biol, vol. 51, no. 6, pp. 749–84, 1989.

[44] S. Nikonov, N. Engheta, and E. Pugh Jr, “Kinetics of recovery of the dark-adapted salamander rod photoresponse.,” J Gen Physiol., vol. 111, pp. 7–37, 1998.

[45] G. Caruso, H. Khanal, V. Alexiadis, F. Rieke, H. Hamm, and E. DiBenedetto, “Mathematical and computational modelling of spatio-temporal signalling in rod phototransduction,” IEE Proc. Syst. Biol., vol. 152, no. 3, pp. 119–137, 2005.

[46] O. Gross, E. N. Pugh, and M. Burns, “Spatiotemporal cgmp dynamics in living mouse rods.,” Biophys. J., vol. 102, no. 8, pp. 1775–1784, 2012.

[47] O. Gross, E. N. Pugh, and M. Burns, “Calcium feedback to cgmp synthesis strongly attenuates single-photon responses driven by long rhodopsin lifetimes.,” Neuron, vol. 76, no. 2, pp. 370–82, 2012.

[48] J. Reingruber, J. Pahlberg, M. Woodruff, A. Sampath, G. Fain, and D. Holcman, “Detection of single photons by rod photoreceptors,” Proc Natl Acad Sci U S A, vol. 110, no. 48, pp. 19378–83, 2013.

[49] T. D. Lamb and T. W. Kraft, “Quantitative modeling of the molecular steps underlying shut-off of rhodopsin activity in rod phototransduction,” Molecular vision, vol. 22, p. 674, 2016.

[50] T. D. Lamb and T. W. Kraft, “A quantitative account of mammalian rod phototransduction with pde6 dimeric activation: responses to bright flashes,” Open biology, vol. 10, no. 1, p. 190241, 2020.

[51] H. R. Matthews and G. L. Fain, “The effect of light on outer segment calcium in salamander rods,” The Journal of Physiology, vol. 552, no. 3, pp. 763–776, 2003.

[52] A. Mendez, M. Burns, I. Sokal, A. Dizhoor, W. Baehr, K. Palczewski, D. Baylor, and J. Chen, “Role of guanylate cyclase-activating proteins (gcaps) in setting the flash sensitivity of rod photoreceptors.,” Proc Natl Acad Sci U S A, vol. 98, pp. 9948–9953, 2001.

[53] V. A. Klenchin, P. D. Calvert, and M. D. Bownds, “Inhibition of rhodopsin kinase by recoverin: further evidence for a negative feedback system in phototransduction,” Journal of Biological Chemistry, vol. 270, no. 27, pp. 16147–16152, 1995.

[54] C.-K. Chen, J. Inglese, R. J. Lefkowitz, and J. B. Hurley, “Ca2+-dependent interaction of recoverin with rhodopsin kinase,” Journal of Biological Chemistry, vol. 270, no. 30, pp. 18060– 18066, 1995.

[55] S. Kawamura and S. Tachibanaki, “Molecular basis of rod and cone differences,” Progress in retinal and eye research, p. 101040, 2021.

[56] S. Kawamura, O. Kuwata, M. Yamada, S. Matsuda, O. Hisatomi, and F. Tokunaga, “Photoreceptor protein s26, a cone homologue of s-modulin in frog retina,” Journal of Biological Chemistry, vol. 271, no. 35, pp. 21359–21364, 1996.

[57] R. D. Hamer and C. W. Tyler, “Phototransduction: Modeling the primate cone flash response,” Visual Neuroscience, vol. 12, no. 6, pp. 1063–1082, 1995.

[58] A. Moriondo and G. Rispoli, “A step-by-step model of phototransduction cascade shows that ca2+ regulation of guanylate cyclase accounts only for short-term changes of photoresponse,” Photochemical and Photobiological Sciences, vol. 2, no. 12, pp. 1292–1298, 2003.

[59] J. M. Angueyra, J. Baudin, G. W. Schwartz, and F. Rieke, “Predicting and manipulating cone responses to naturalistic inputs,” Journal of Neuroscience, vol. 42, no. 7, pp. 1254–1274, 2022.

[60] Y. Li, F. Falleroni, S. Mortal, U. Bocchero, D. Cojoc, and V. Torre, “Calcium flares and compartmentalization in rod photoreceptors,” Proceedings of the National Academy of Sciences, vol. 117, no. 35, pp. 21701–21710, 2020.

[61] N. Ingram, A. Sampath, and G. Fain, “Voltage-clamp recordings of light responses from wt and mutant mouse cone photoreceptors.,” J Gen Physiol, vol. 151, no. 11, pp. 1287–1299, 2019.

[62] C. J. Beelen, S. Asteriti, L. Cangiano, K.-W. Koch, and D. Dell’Orco, “A hybrid stochastic/deterministic model of single photon response and light adaptation in mouse rods,” Computational and Structural Biotechnology Journal, vol. 19, pp. 3720–3734, 2021.

[63] R. Hamer, S. Nicholas, D. Tranchina, T. Lamb, and J. Jarvinen, “Toward a unified model of vertebrate rod phototransduction,” Vis. Neurosci., vol. 22, pp. 417–436, 2005.

[64] S. Nikonov, T. Lamb, and E. Pugh Jr, “The role of steady phosphodiesterase activity in the kinetics and sensitivity of the light-adapted salamander rod photoresponse,” J. Gen. Physiol., vol. 116, pp. 795–824, 2000.

[65] C. Klaus, G. Caruso, V. V. Gurevich, H. E. Hamm, C. L. Makino, and E. DiBenedetto, “Phototransduction in retinal cones: Analysis of parameter importance,” PloS one, vol. 16, no. 10, p. e0258721, 2021.

[66] P. Bisegna, G. Caruso, D. Andreucci, L. Shen, V. Gurevich, H. Hamm, and E. DiBenedetto, “Diffusion of the second messengers in the cytoplasm acts as a variability suppressor of the single photon response in vertebrate phototransduction,” Biophys. J., vol. 94, pp. 3363–3383, 2008.

[67] E. Pugh Jr and T. Lamb, “A quantitative account of the activation steps involved in phototransduction in amphibian photoreceptors,” J. Physiol., vol. 449, pp. 719–758, 1992.

[68] M. Mazzolini, G. Facchetti, L. Andolfi, R. P. Zaccaria, S. Tuccio, J. Treu, C. Altafini, E. M. Di Fabrizio, M. Lazzarino, G. Rapp, et al., “The phototransduction machinery in the rod outer segment has a strong efficacy gradient,” Proceedings of the National Academy of Sciences, vol. 112, no. 20, pp. E2715–E2724, 2015.

[69] J. Reisert and J. Reingruber, “Ca2+-activated clcurrent ensures robust and reliable signal amplification in vertebrate olfactory receptor neurons.,” Proc Natl Acad Sci U S A, vol. 116, no. 3, pp. 1053–1058, 2019.

[70] I. Peshenko, E. Olshevskaya, A. Savchenko, S. Karan, P. K W. Baehr, and A. Dizhoor, “Enzymatic properties and regulation of the native isozymes of retinal membrane guanylyl cyclase (retgc) from mouse photoreceptors.,” Biochemistry, vol. 50, no. 25, pp. 5590–600, 2011.

[71] I. Peshenko and A. Dizhoor, “Ca2+ and mg2+ binding properties of gcap-1. evidence that mg2+-bound form is the physiological activator of photoreceptor guanylyl cyclase.,” J Biol Chem., vol. 281, no. 33, pp. 23830–41, 2006.

[72] T. Ohyama, D. H. Hackos, S. Frings, V. Hagen, U. B. Kaupp, and J. I. Korenbrot, “Fraction of the dark current carried by ca2+ through cgmp-gated ion channels of intact rod and cone photoreceptors,” The Journal of general physiology, vol. 116, no. 6, pp. 735–754, 2000.

[73] S. Nikonov, R. Kholodenko, J. Lem, and E. Pugh Jr, “Physiological features of the s- and m-cone photoreceptors of wild-type mice from single-cell recordings.,” J Gen Physiol, vol. 127, no. 4, pp. 359–74, 2006.

[74] N. Ingram, A. Sampath, and G. Fain, “Why are rods more sensitive than cones?,” J Physiol, vol. 594, no. 19, pp. 5415–26, 2016.

